# Complex coacervation reshapes the aggregation landscape of tau

**DOI:** 10.64898/2026.06.27.734011

**Authors:** Zexiang Han, Peifeng Xu, Yangteng Ou, Daoyuan Qian, Zhenting Xiao, Yuxiang Wu, Alessia Santambrogio, Michele Vendruscolo, Tuomas P. J. Knowles

## Abstract

Biomolecular condensates are increasingly implicated in protein aggregation, yet their contribution is often reduced to that of concentrating reactants. Whether the phase state of a protein itself changes how it aggregates remains unclear. Using complex coacervates of a tau repeat-domain construct (K12) with heparin, we show that phase separation does not simply accelerate tau aggregation but redirects it along a distinct kinetic regime. Amyloid nucleation and growth are largely associated with the condensed phase, where intact phase-separated mixtures nucleate with a markedly shortened lag phase, whereas the corresponding dilute phase contributes little to overall amyloid formation. Aggregation kinetics in this regime become largely decoupled from total protein concentration. Because phase equilibrium pins the composition of the dense phase, additional tau partitions largely into the coexisting dilute phase without substantially altering the reacting population. This weak concentration dependence provides a kinetic signature of compartmentalized aggregation, and it recurs across chemically distinct coacervates formed with heparin, RNA, and polyglutamate, pointing to a general feature of coacervate-mediated tau assembly rather than a heparin-specific effect. Aggregation within coacervates also yields fibrils with altered morphology and secondary structure, suggesting access to alternative regions of the assembly landscape, and shows reduced sensitivity to bulk pH perturbations. Together, these results show that condensation changes tau aggregation by defining the local reaction environment: phase equilibrium buffers the dense-phase composition, in turn altering aggregation kinetics and the properties of the amyloid formed.

## Introduction

Protein misfolding and aggregation are characteristic of many neurodegenerative conditions, including Alzheimer’s disease and Parkinson’s disease.^1–3^ The microtubule-associated protein tau is central to a group of such disorders known as tauopathies, and tau aggregation into neurofibrillary tangles correlates with cognitive decline and neuronal loss.^4–7^ Previous studies of amyloid formation have provided a fundamental understanding of this aggregation process.^8–11^ However, the recent discovery that tau and other amyloid-associated proteins undergo liquid-liquid phase separation (LLPS) to form biomolecular condensates has shifted our view of this pathological cascade.^12–17^ A central question is how these condensed entities, which are hubs of cellular activity, are linked to pathogenic aggregation.^18–20^

The prevailing hypothesis has been that condensates accelerate the aggregation process primarily by achieving a very high concentration of proteins within the dense phase. Many such condensates undergo liquid-to-solid transition, accompanied by conformational rearrangements of the proteins and characterized by a loss in internal molecular dynamics over time, eventually maturing into amyloid-like structures.^13,14,21,22^ High-resolution imaging by total internal reflection fluorescence (TIRF) or transmission electron microscopy (TEM) further shows that amyloid fibrillation can occur within or at the interface of condensates, supporting the view that these dense liquid phases can function as crucibles for amyloid formation.^15,23–25^

Nevertheless, biomolecular condensates are not passive concentrators. They establish unique physicochemical microenvironments that differ from the surrounding dilute phase.^26,27^ Key properties such as the presentation of a dilute-dense phase interface, altered microviscosity, intermolecular interaction potentials, and distinct chemical interior could profoundly influence the protein aggregation kinetics and pathways.^24,28–31^ Existing condensates can actively modulate α-synuclein aggregation, either accelerating it when amyloidogenic proteins localize at interfaces or suppressing it through sequestration, indicating that the outcome is dictated by the nature of the physical interaction.^32^ In homotypic prion-like domain systems, condensate interiors have been shown to suppress fibril formation by acting as sinks, with nucleation instead localized to condensate interfaces^30^, illustrating that the dense phase is not uniformly aggregation-promoting. Although recent work has begun to clarify the connection between condensation and aggregation, our quantitative and mechanistic understanding of the process remains incomplete.^8,33^ Specifically, whether condensates alter the fibrillation kinetics, produce structurally distinct fibrils, or modulate how aggregation responds to environmental cues remains unresolved.

In this study, we use tau-heparin mixtures to ask how complex coacervation shapes the aggregation landscape of tau. We find that condensation does not simply accelerate amyloid aggregation relative to bulk solution but redirects it along an alternative pathway with a distinct kinetic signature. Aggregation within coacervates becomes largely decoupled from total protein concentration, a consequence of the dense-phase composition being constrained by phase equilibrium. This compartmentalized regime recurs across chemically distinct polyanions and, as a functional consequence, renders aggregation markedly less sensitive to bulk pH perturbations. Taken together, these results establish the phase state of tau as a determinant of both the kinetics and the products of its amyloid assembly.

## Results

### Tau liquid-liquid phase separates with heparin

Tau does not readily form aggregates on its own, making cofactors widely used ingredients for in vitro aggregation and diagnostic assays.^34–36^ We establish a model system using the K12 tau fragment (hereafter referred to as tau), which spans the three microtubule-binding repeat regions that form the ordered filament cores of tau fibrils in both AD and Pick’s disease (**Supplementary Figure S1**).^37^ Using heparin as a cofactor, we employ this system to dissect how phase state influences amyloid aggregation. **Figure 1** provides a characterization of the phase behavior of tau-heparin mixtures.

**Figure 1.**
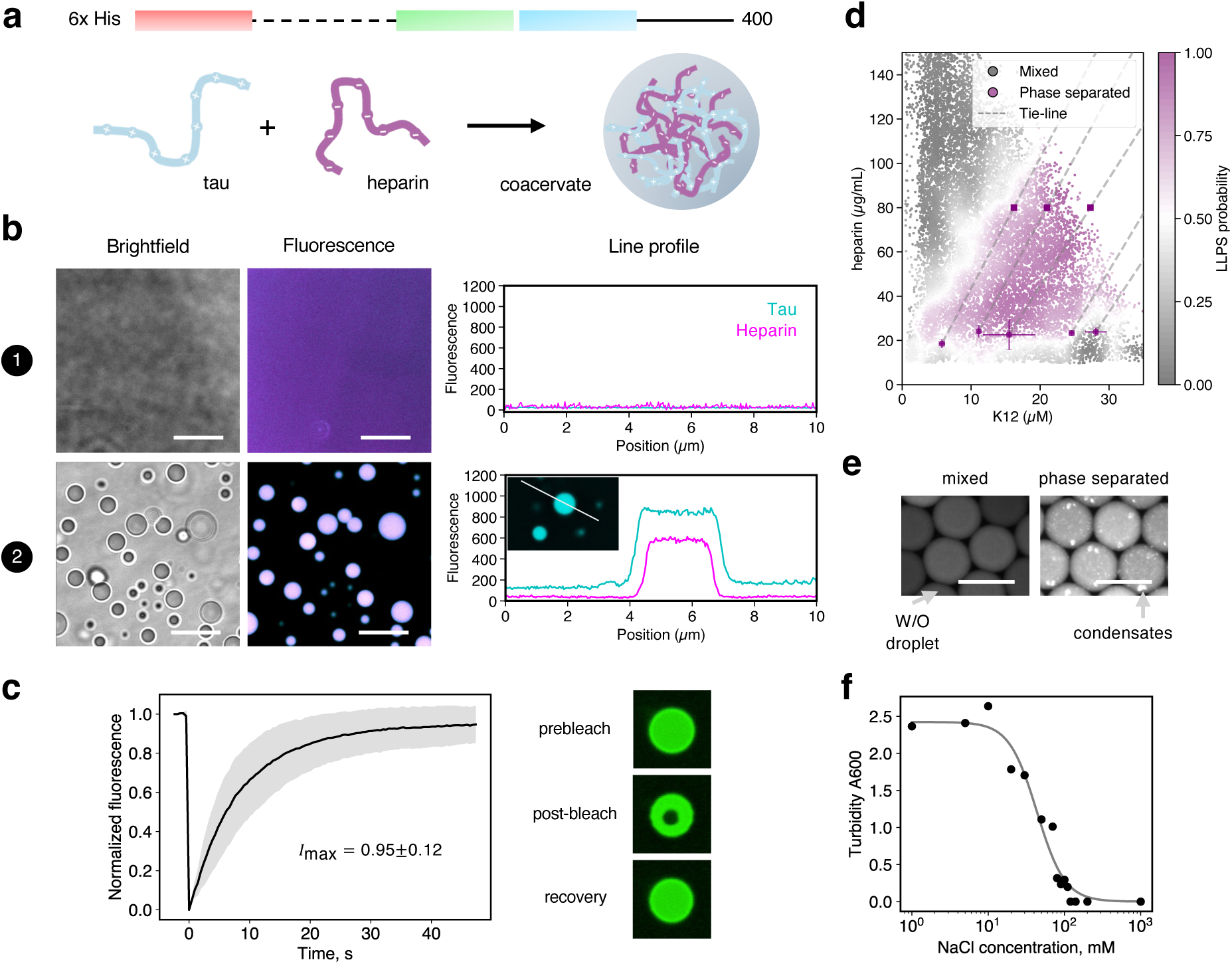
Tau and heparin form liquid-like condensates by complex coacervation. (a) Schemes of tau fragment sequence and tau-heparin LLPS by complex coacervation. (b) Brightfield and confocal images of tau-heparin mixtures prepared at: (1) 5 µM tau, 80 µg mL^−1^ heparin; (2) 60 µM tau, 80 µg mL^−1^ heparin. Both biomolecules are covalently tagged with fluorescent labels: tau (cyan) and heparin (magenta). Shown on the right are representative fluorescence line profiles. Scale bars = 10 µm. (c) FRAP assay: normalized fluorescence recovery kinetics of freshly prepared condensates (*n* = 3) and representative FRAP images. (d) High-resolution phase diagram measured using droplet microfluidics (number of data points *n* = 49,522). Approximate tie line proxies were inferred by connecting each parent composition to its corresponding dilute-phase composition. (e) Fluorescence images of water-in-oil (W/O) microdroplets containing tau protein in mixed and phase-separated states. Scale bars = 100 µm. (f) Turbidity assay using absorbance measurement at 600 nm, showing the dissolution of condensates with increasing NaCl addition. Condensates are prepared using 20 µM tau and 80 µg mL^−1^ heparin. Gray line represents the sigmoidal fit of the data.

Within some concentration thresholds, tau protein and heparin phase separate into high-circularity, biomolecule-dense droplets surrounded by a dilute phase (**Figure 1**b). Both biomolecules colocalize within the condensates, indicative of associative phase separation. This is mediated by electrostatic attraction between negatively-charged heparin and the microtubule-binding domain of tau that is lysine- and arginine-rich. Fluorescence recovery after photobleaching (FRAP) of condensates reveals their highly liquid-like nature, with near-complete (95%) recovery of the initial fluorescence intensity (**Figure 1**c).

We mapped out the tau-heparin phase diagram at ultrahigh resolution using combinatorial droplet microfluidics^38,39^, revealing the precise phase boundary (**Figure 1**d, e). The system shows re-entrant behavior characteristic of complex coacervation, a main mode of phase separation for biomolecules^40^: for instance, for 20 µM protein, the mixture transitions from a soluble state to a phase-separated state before transitioning back to a soluble state with increasing heparin concentration, as separately verified using turbidity measurements (**Supplementary Figure S2**). This re-entrance arises from charge and/or valency balancing, where phase separation occurs around charge stoichiometry, but excess heparin results in condensate re-solubilization.^41,42^ Additionally, the tie line slope was found to be positive, reaffirming the associative interaction between tau and heparin within the condensed phase.

Lastly, as an orthogonal validation, we used sodium chloride as a perturbant to study its effect on tau-heparin LLPS driven by associative, charged interactions. Indeed, the turbidity of the mixture decreased progressively with increasing NaCl concentration and was fully abolished above the critical salt concentration of approximately 120 mM, yielding a homogeneous solution phase due to electrostatic screening of biomolecular charges by salt ions (**Figure 1**f).

### Phase state defines distinct aggregation reaction environments

We explore the compositional phase space of tau-heparin mixtures to investigate how tau condensation influences aggregation. At a constant heparin concentration, systematically increasing tau concentration drives the system from a homogeneous solution to a phase-separated state (**Figure 2**a). The evolution of the physical state of the system is corroborated by dynamic light scattering (DLS) measurements showing a progression from small molecular complexes in solution to subcritical, pre-LLPS nanoclusters to micrometer-scale condensates (**Figure 2**b). This setup allows us to study aggregation in condensates formed purely by protein and polyanionic cofactor without introducing exogenous molecules such as crowding agents that could confound the aggregation process.

**Figure 2.**
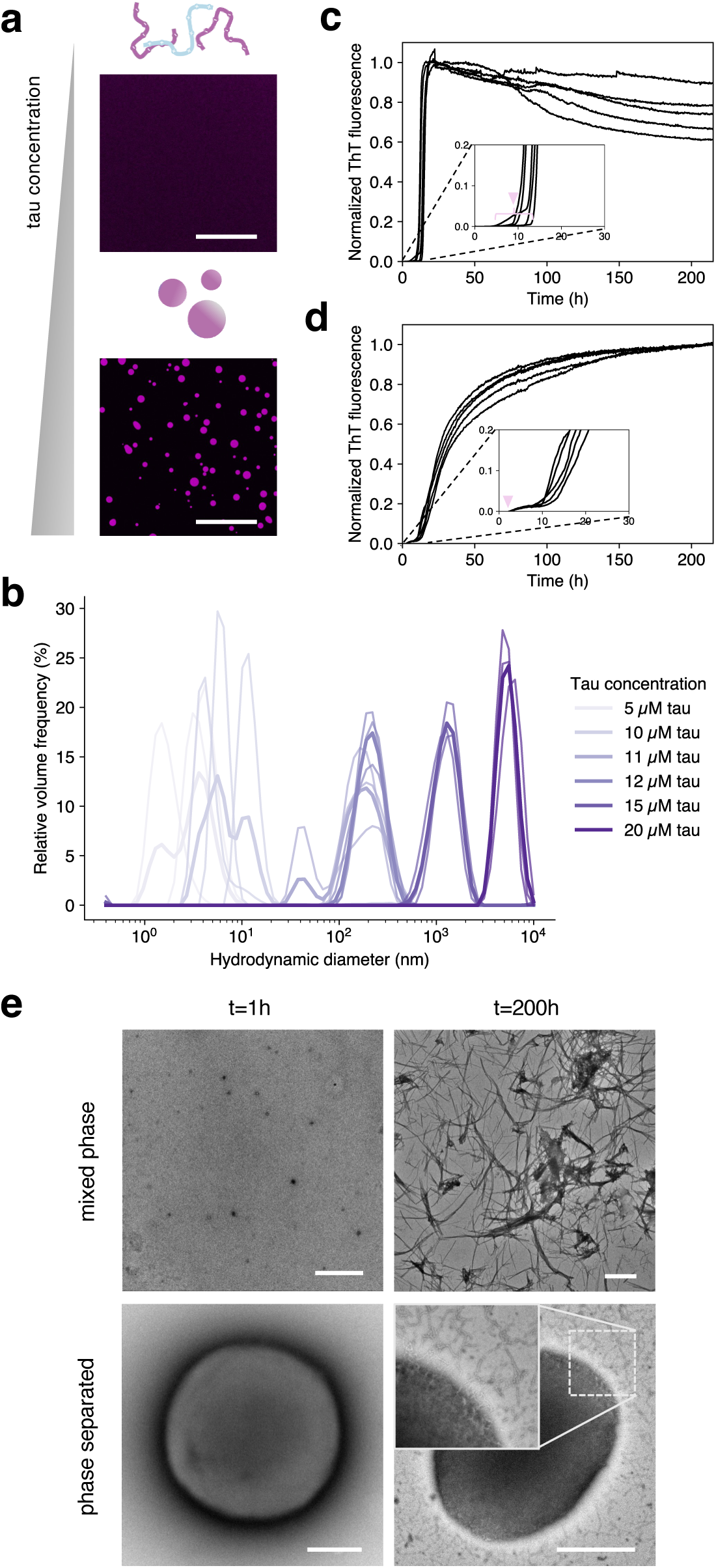
Phase separation alters the kinetics and spatial localization of tau aggregation. (a) Scheme and confocal images of tau-heparin mixtures. Top: mixed phase containing 9.6 µM tau and 80 µg mL^−1^ heparin. Bottom: phase-separated mixture containing 35.5 µM tau and 80 µg mL^−1^ heparin. Scale bars = 50 µm. (b) DLS size distribution of tau-heparin mixtures. Shown are three independent repeats and their average in thicker lines. Samples were prepared in the presence of 80 µg mL^−1^ heparin. (c, d) Corresponding ThT kinetic traces of tau in (c) mixed solution and (d) phase-separated mixture. Reactions were performed in quintuplicates. (e) TEM images of initial mixture and aggregated products from (top) mixed solution and (bottom) phase-separated mixture. Scale bars = 1 µm.

The temporal dynamics of tau amyloid fibrillation in homogeneous and phase-separated reaction environments were monitored using Thioflavin T (ThT) as an amyloid-reporting dye. The normalized ThT kinetic profiles differ qualitatively for samples in the two phase states (**Figure 2**c, d). In the mixed phase, the aggregation curves are archetypically sigmoidal with a steep incline and reach a saturation plateau. Conversely, the traces in the LLPS regime feature a continually increasing ThT signal, indicative of a more dynamic amyloid formation process. Zoomed-in view showing minimal lag phase in phase separated mixtures indicates that nucleation is accelerated. TEM imaging confirms the formation of fibrillar aggregates under both homogeneous and phase-separated conditions (**Figure 2**e). Notably, in phase-separated systems, fibrils are associated with condensate-derived material, suggesting that aggregation occurs in the condensed phase. While these observations establish that phase state alters both aggregation kinetics and the spatial distribution of amyloid products, they do not resolve how individual steps of the aggregation process are affected.

We therefore address the nucleation and growth processes in a semi-decoupled fashion (**Figure 3**a). First, to determine the role of condensates in nucleation, we compared the aggregation kinetics of the parent phase-separated mixture containing 35.5 µM tau against the corresponding isolated dilute phase extracted by centrifugation (**Figure 3**b). Specifically, we examined the lag time *t*_lag_ to quantify how the condensed phase modulates the onset of aggregation. Since the lag phase of amyloid formation integrates multiple microscopic kinetic events in addition to nucleation, different orbital shaking speeds were employed to differentially amplify specific pathways.^43^ Under quiescent conditions, primary and secondary nucleation govern the process, whereas introduction of mechanical agitation by shear promotes fibril fragmentation.

**Figure 3.**
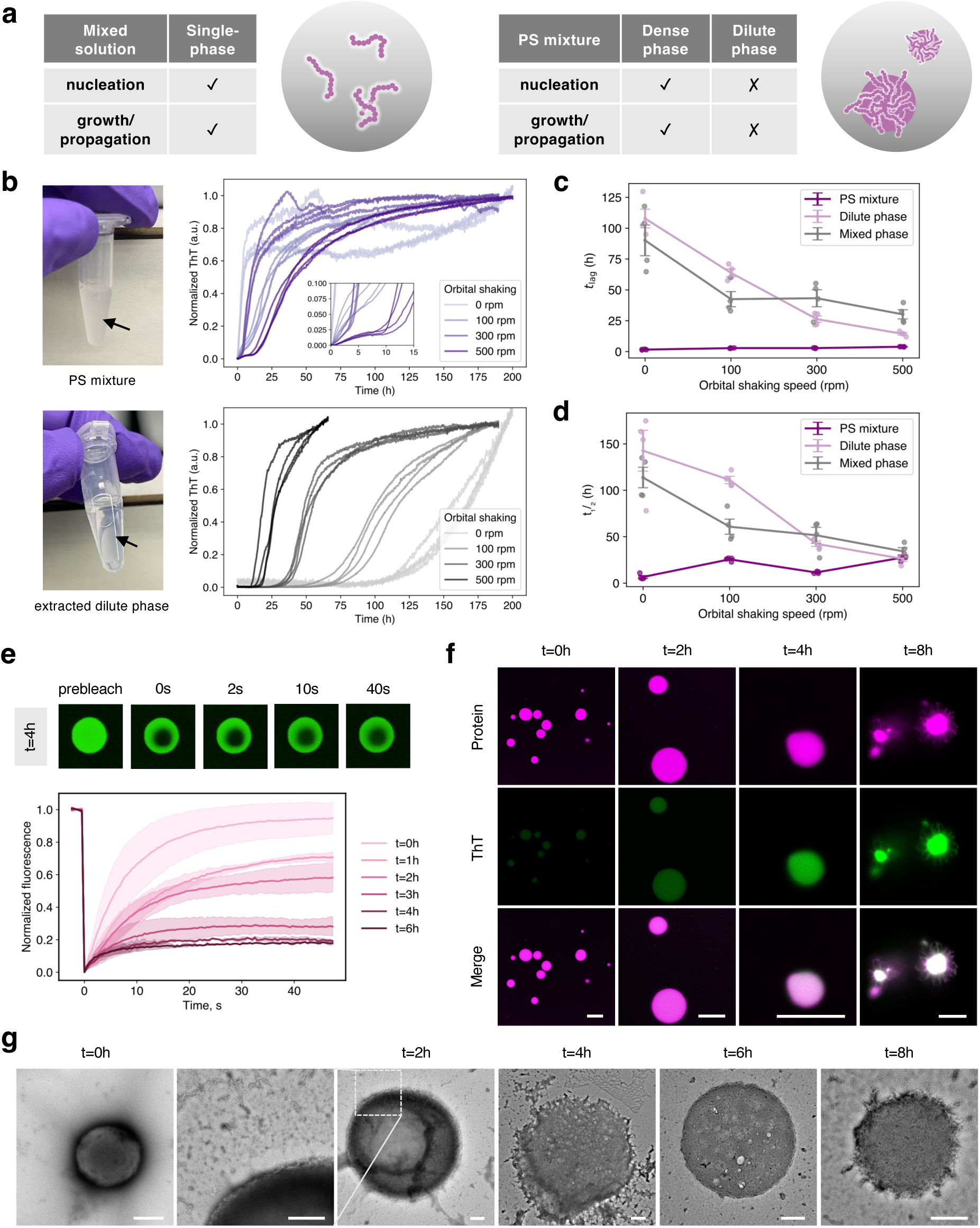
Tau amyloid nucleation and growth are associated with the condensate rather than the bulk dilute phase. (a) Tables and schemes summarizing the compartmentalized nature of nucleation and growth of amyloid fibrillation in different phase states. (b) Photographs and ThT aggregation kinetic traces for phase-separated mixtures and their corresponding dilute phases. The phase-separated mixture contained 35.5 µM tau and 80 µg mL^−1^ heparin. Reactions were performed under different orbital shaking regimes in quadruplicates. (c, d) Effect of shear on aggregation (c) lag time *t*_lag_ and (d) half-time *t*_1/2_ for phase-separated mixtures, corresponding dilute-phase samples, and mixed phase samples. Mixed phase samples contained 7.0 µM tau and 80 µg mL^−1^ heparin. (e) Normalized FRAP curves demonstrating liquid-to-solid transition. Shaded are error bands. Shown above are representative images showing FRAP recovery in aged condensates. (f) Confocal images of condensates following different incubation periods. Tau proteins were labeled with Alexa Fluor 546 NHS ester (magenta) and amyloids stained using ThT (green). Scale bars indicate from left to right 10 µm, 10 µm, 5 µm, and 10 µm. Incubation was performed under identical conditions as microplate-based aggregation assay. (g) TEM images illustrating the progressing ageing of tau condensates with increased amount of fibrillar network inside. Scale bars = 500 nm.

Both homogeneous (with a tau concentration of 7 µM) and isolated dilute-phase samples exhibit shear-dependent nucleation and growth as quantified by *t*_lag_ and aggregation half-time *t*_1/2_ (**Figure 3**c, d). Intact phase-separated samples, however, aggregated robustly under all conditions, showing nearly spontaneous nucleation with an averaged lag phase of 2.8 h (as indicated per **Figure 2**d and **Figure 3**b insets). This is in stark contrast to the isolated coexisting dilute phase, which only aggregated after ∼108 h of incubation under quiescent condition. This disparity suggests that although the dilute phase maintains a latent ability to nucleate under high-shear conditions, the condensed phase provides a more permissive microenvironment that lowers the kinetic barrier to nucleate amyloid formation.

To resolve the spatial-temporal progression of amyloid growth, we combined confocal microscopy, FRAP assay and TEM imaging. We show that the slow and persistent increase in ThT fluorescence measurements in bulk is attributed to the gradual maturation of tau coacervates into solid-like, amyloid assemblies, reflecting amyloid growth inside condensates (**Figure 3**e, f). Tau condensates undergo a progressive liquid-to-solid transition with prolonged incubation, as evidenced by a time-dependent decrease in the mobile fraction, indicating an increasingly rigid material state (**Figure 3**e). This loss of condensate fluidity is accompanied by a concurrent rise in ThT fluorescence, with ThT signal colocalizing with that of the condensate phase (**Figure 3**f). This suggests that the formation of β-sheet-rich fibrils occurs within the condensed phase. TEM imaging at discrete ageing time points reveals an increasing density of fibrillar aggregates within condensates, supporting this interpretation (**Figure 3**g). Over time, fibrils elongate into the dilute phase. The bulk ThT signal continued to increase while intra-condensate signal plateaued, reflecting the saturation of ThT-binding sites within the condensate volume and consistent with the observation of fibril elongation from condensate interface into the dilute phase (**Supplementary Figure S3**). In contrast to the long, parallel fibril bundles typically observed in bulk solution, the condensates eventually transform into a dense, percolated fibrillar amyloid network that permeates the droplet volume, with flexible fibrils extending across the interface (**Figure 2**e).

Finally, comparison of the relative contribution to the final ThT signal between phase-separated samples and isolated dilute phases showed that the dilute phase contributes a significantly smaller fraction of the total fluorescence intensity, even under the highest applied shear stress (**Supplementary Figure S4**). These observations further demonstrate that the bulk ThT signal originates primarily from the condensed phases.

### Liquid phase condensation alters the tau aggregation kinetic pathway

Having established that amyloid aggregation is largely taking place within condensates and that bulk fluorescence can faithfully report this internal maturation process, we next sought to quantify how the phase state itself impacts the amyloid aggregation behavior. Systematic mapping of aggregation half-times *t*_1/2_ across tau-heparin compositions reveals that phase separation reshapes the aggregation kinetic landscape (**Figure 4**a), with the dashed line marking the transition between homogeneous and macroscopically phase-separated regimes. To explore how phase state modulates the aggregation pathway in detail, we probed aggregation kinetics for a tau dilution series at three different constant heparin concentrations, each driving the system from a mixed solution to a phase-separated mixture (**Figure 4**b). The monomer concentration dependence of *t*_1/2_ reveals the two distinct regimes that map directly onto the physical state of the system per DLS measurements (**Figure 2**b).

**Figure 4.**
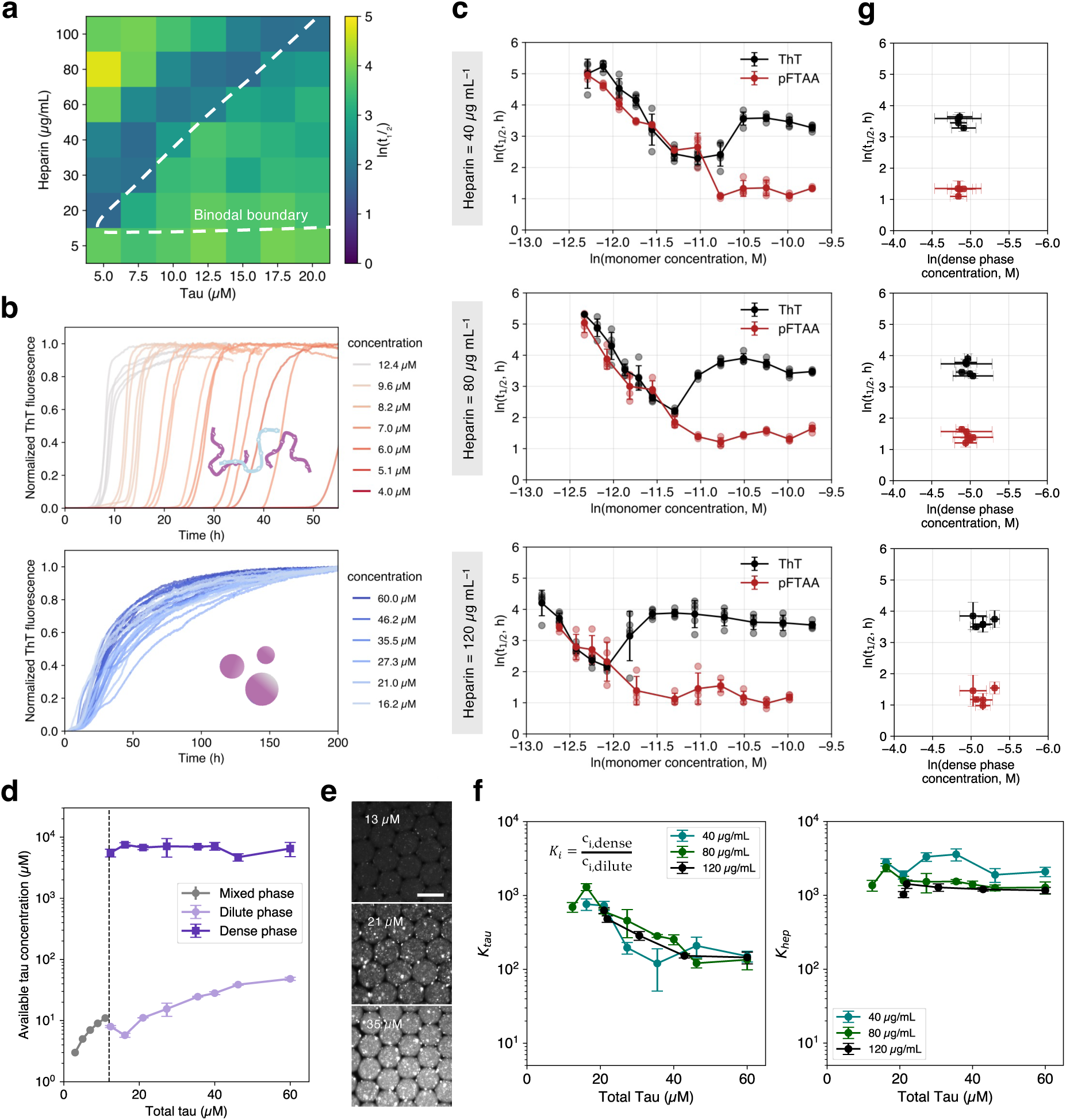
Phase separation decouples tau aggregation kinetics from total protein concentration. (a) Aggregation half-time *t*_1/2_ of tau-heparin mixtures across compositional and phase space, illustrating phase state-dependent amyloid formation behavior. Dashed line delineates the phase boundary between mixed and phase-separated states. (b) Normalized ThT kinetic traces for a dilution series of tau in the presence of 80 µg mL^−1^ heparin: (top) as a homogeneous solution, (bottom) as a phase-separated mixture. (c) Relationship between tau monomer concentration and aggregation half-time *t*_1/2_ in the presence of 40, 80 and 120 µg mL^−1^ heparin, respectively. Aggregation assay was performed in quintuplicates. (d) Measured tau concentrations in the presence of 80 µg mL^−1^ heparin in different phases. Zoomed-in views show the relationship between the reaction-relevant tau concentrations and total tau concentrations. (e) Fluorescence images of tau-heparin condensates formed in water-in-oil microdroplets. Images show increasing total tau concentration from top to bottom in the presence of 80 µg mL^−1^ heparin, where the dilute phase becomes significantly more enriched in tau that is fluorescently labeled. Scale bars = 100 µm. (f) Biomolecular partitioning behavior as quantified by partition coefficients across coacervate compositions. (g) Rescaling ln(*t*_1/2_) against dense phase tau concentration in LLPS regime.

In the mixed phase, *t*_1/2_ exhibits the strong dependence on tau concentration that is characteristic of a nucleation-polymerization reaction. Increasing tau monomer concentration leads to faster nucleation and growth. This is expected from classical aggregation models where higher monomer availability fuels secondary nucleation, confirmed by global kinetic fitting (**Supplementary Figure S5**).^44,45^ We evaluated the power-law scaling exponent describing how the aggregation half-time scales with total tau concentration (**Supplementary Figure S6**). An apparent scaling exponent of approximately −3 was observed in mixed solutions, indicating a strong dependence of aggregation kinetics on concentration. Consequently, a modest increase in total tau concentration leads to substantial reductions in aggregation half-time, consistent with rapid fibrillation in the mixed phase.

Strikingly, upon entering the LLPS regime, the monomer concentration dependency is drastically reduced. This flattening of concentration dependence was observed across multiple heparin concentrations, with the transition point shifting with the corresponding binodal positions (**Figure 4**c). This highlights a phase state-specific shift in the aggregation kinetic landscape.

In addition to ThT that reports on the formation of well-ordered, canonical amyloid structures, an oligothiophene dye pFTAA was used to detect early-stage intermediates, providing complementary insights into the aggregation pathway (**Supplementary Figure S7**).^46^ Shown in **Figure 4**c, ThT aggregation kinetics exhibited a non-monotonic concentration dependence from mixed to phase-separated states: aggregation half-time *t_1/2_* initially decreased with increasing tau concentration, then transiently increased at intermediate concentrations before resuming a decrease at higher concentrations. This abrupt change in half-time suggest a competition between distinct assembly routes in the case of tau nanoclusters, sized around 200 nm, that bridge the two macroscopic phase states. In contrast, the pFTAA-based t_1/2_ decreased consistently and then levelled off upon entry into the LLPS regime, plateauing within the first 6 h in phase-separated mixtures, well before the rise of the ThT signal, whose t_1/2_ saturates at 40 ± 9 h under the same conditions. These data support the view that condensates accelerate the initial nucleation step, enabling rapid formation and accumulation of prefibrillar species, whereas the subsequent conversion of these species into more canonical-like amyloid fibrils proceeds more slowly within condensates.

To rationalize the negligible dependence of *t_1/2_* on total tau concentration in the phase-separated regime, we consider the role of biomolecular and phase partitioning in defining the effective reaction environment (**Supplementary Figure S8**). In tau-heparin coacervates, phase equilibrium constrains the compositions of coexisting dense and dilute phases (**Figure 4**d). For a given heparin concentration, increasing tau concentrations above the binodal results in elevated dilute-phase tau concentration and increased volume fractions, whereas the dense phase tau concentration remains relatively constant (6.5 ± 0.8 mM) over a broad range of total tau concentrations (**Supplementary Figure S9**). Above the binodal that defines the saturation concentration, excess tau species partition preferentially into the dilute phase (**Figure 4**e, f). Therefore, the dense phase represents a compositionally defined coacervate state constrained by the tau-heparin electrostatic attraction, with an electrostatically defined tau-to-heparin stoichiometry in the condensate phase estimated to be 3.0 ± 0.6. This saturated dense-phase limit is typical of complex coacervates formed between polyelectrolytes.^47,48^

Consistent with this framework, the minimal dependence of aggregation kinetics on total tau concentration arises from a buffered dense phase across the explored conditions. Under these constraints, increases in total tau primarily do not substantially alter the composition of the condensed phase in which nucleation and growth occur, and the dense phase chemical potential is effectively pinned. The invariance of kinetics with respect to additional tau entering the dilute phase further supports the conclusion that the dilute-phase reservoir does not contribute measurably to the aggregation pathway under these conditions (**Supplementary Figure S10**). Consequently, the resulting data collapse into a cluster of points corresponding to the nearly invariant dense-phase composition and the associated *t_1/2_*values (**Figure 4**g). The absence of a clear monomer scaling thus represents a kinetic signature of compartmentalized aggregation, in which fibrillation proceeds within condensates that are compositionally defined through phase equilibria.

Together, these results show that the apparent concentration dependence of tau aggregation is modulated by phase separation. Given that the coexisting dilute phase does not meaningfully contribute to amyloid fibrillation, aggregation kinetics in the phase-separated regime are governed by the intra-condensate chemical potential that defines the aggregation microenvironment, thereby bridging thermodynamic phase behavior with aggregation kinetics.

### Phase state-dependent amyloidogenesis can yield distinct amyloid states

While kinetic differences could in principle arise from a single pathway with altered rate constants, differences in the final product would indicate that condensation selects for an alternative free energy minimum on the tau assembly landscape (**Figure 5**a).^49^ TEM images revealing structural differences between the fibrils, in conjunction with the distinct responses of ThT and pFTAA during the same fibrillation process, strongly suggest phase state-dependent amyloid assembly.

**Figure 5.**
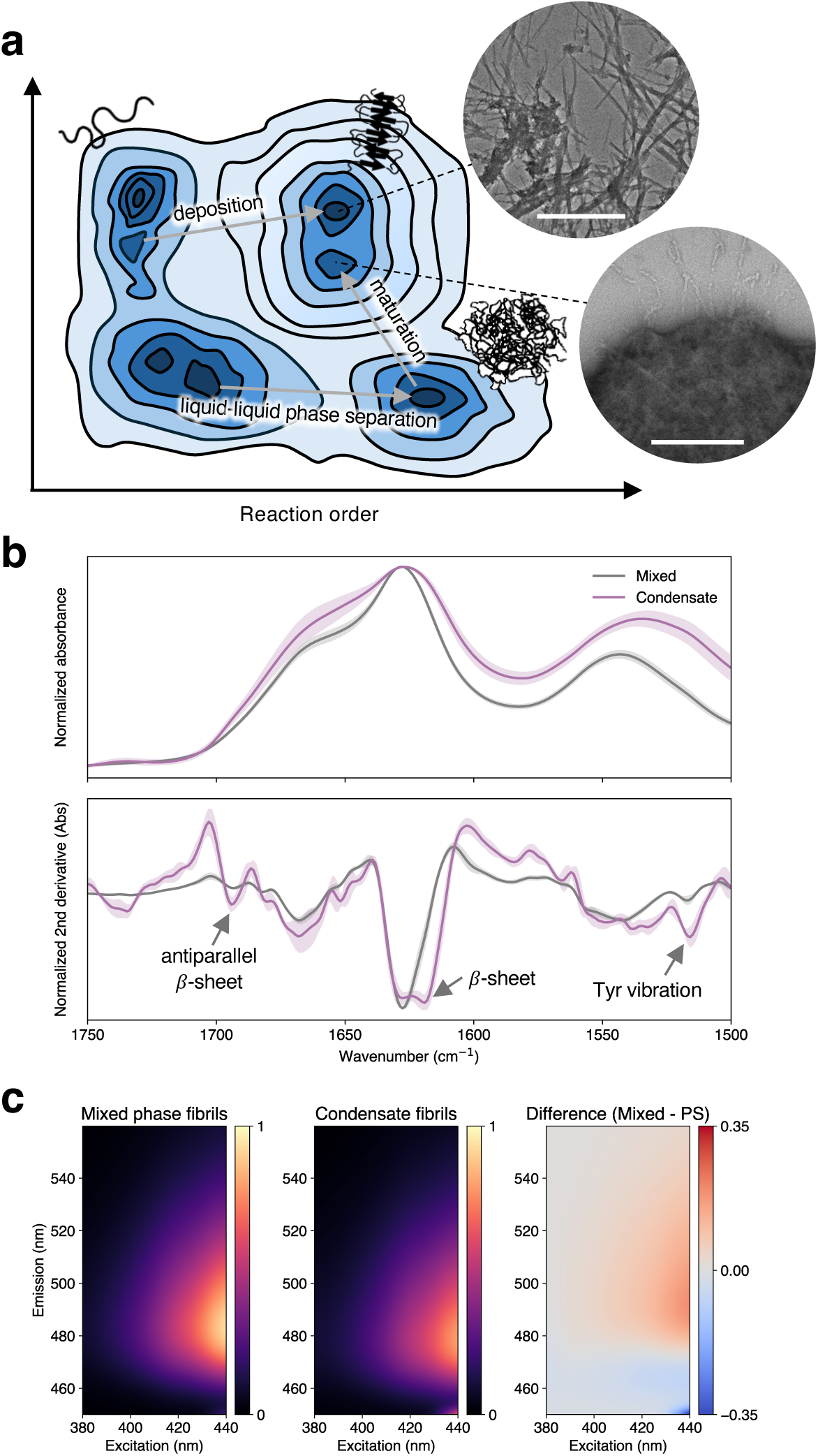
Deposition and maturation pathways lead to distinct amyloid states. (a) Proposed scheme of energy landscape showing the conversion between the native, droplet, and amyloid states of tau proteins. Insets are TEM images of as-formed amyloid fibrils formed from mixed solution or from condensates, illustrating two different amyloid states. Scale bars = 1 µm (top) and 500 nm (bottom). (b) Averaged FTIR spectra and second derivative plot showing the spectral fingerprints of fibrils grown in mixed, homogeneous solutions versus in condensates. Error bands show the standard deviations. (n=4) (c) Normalized 2D excitation-emission maps for mixed-phase and phase-separated samples with ThT as amyloid-binding dye. Shown on the right is the calculated difference between the two plots, highlighting the spectral shift in emission peak.

Fourier-transform infrared (FTIR) spectroscopy was used to probe secondary structure content of the as-grown fibrils. In the second derivative plot, the amide I band (1600–1700 cm⁻¹) of fibrils from mixed solution exhibited a minimum at ∼1628 cm⁻¹, characteristic of β-sheet structure typical of amyloid deposits (**Figure 5**b). On the other hand, fibrils formed within the condensate phase maintained the β-sheet signature but displayed a shifted and split-line shape, accompanied by increased contribution of antiparallel β-sheet organization (**Supplementary Figure S11**).^50^ This points to the formation of different fibril polymorphs, with additional spectral differences reflecting an altered hydrogen-bonding network and hierarchical assembly associated with the dense-phase environment. Furthermore, the spectral properties of ThT can report on the local environment of bound fibrils.^51^ For amyloid fibrils in homogeneous solutions, the 2D excitation-emission heatmap displayed a characteristic excitation/emission maximum at ∼440/481 nm, whereas fibrils formed in condensates exhibited a blue-shifted emission maximum at ∼475 nm (**Figure 5**c and **Supplementary Figure S12**). The difference map clearly visualizes this spectral shift, indicating that ThT experiences a distinct binding environment on condensate-derived fibrils.

### Generality of phase state-dependent amyloid assembly

Beyond the specific case of heparin, we examined tau coacervation with poly-rU RNA and polyglutamate (**Supplementary Figure S13**) and how phase state affects amyloid fibrillation. These systems span distinct classes of biomacromolecules – including glycosaminoglycans, nucleic acids and polypeptides – that differ in their backbone chemistry, charge distribution, and interaction modes with tau. As such, they provide an informative set of comparisons for assessing phase state-dependent amyloid aggregation.

Despite their distinct chemical identities, which could influence tau binding and conformation^52^, all tested systems exhibit phase state-dependent aggregation kinetics (**Supplementary Figure S14**). In each case, entry into the phase-separated regime is accompanied by a marked reduction in the dependence of aggregation kinetics on total protein concentration, indicating that the governing variable is no longer bulk composition but the properties of the condensed phase. The convergence of this behavior across chemically distinct coacervates suggests that it arises from generic features of the condensate microenvironment, such as constrained composition, crowding and spatial confinement, rather than from specific molecular interactions unique to a given polyanion. The observation that RNA-tau condensates recapitulate this behavior suggests that such change in aggregation may be relevant in cellular contexts, where tau engages in heterotypic interactions with nucleic acids and other biomolecules.^53^ Finally, FTIR spectroscopic analysis of the as-formed fibrils reveals detectable, albeit more modest, differences in the fibril secondary structures (**Supplementary Figure S15**). This indicates that although the structural polymorph selection may remain sensitive to molecular chemistry, the underlying concentration dependence of the aggregation kinetics is consistently phase state-dependent.

Taken together, through multi-scale imaging, spectroscopic and kinetic analyses, we have established phase state as an important parameter in tau amyloid formation. LLPS creates micro-compartmentalized reactors wherein physical confinement and macromolecular crowding impose an amyloid aggregation pathway unlike that observed in homogeneous solutions. Beyond concentrating reactants, condensation does not merely modulate the kinetics of a conserved fibrillation process but can redirect tau along a different molecular assembly trajectory characterized by unique kinetic fingerprints and possibly distinct amyloid structural profiles.

### Functional consequence of phase state-dependent amyloid aggregation

Lastly, because phase separation redefines the effective reaction environment for tau aggregation, we asked whether this altered state affects how aggregation responds to solution perturbations. pH is a key regulator of amyloid formation in homogeneous systems^54,55^, but its influence on fibrillation within condensed phases remains unclear. Here, we show that phase separation also alters the pH dependence of tau amyloid formation, revealing that the dense phase can buffer against environmental perturbation effects on fibrillation.

Biomolecular condensates establish distinct physicochemical microenvironments, such pH gradients^56–58^, that decouple reactions from bulk solution conditions. Using pH-sensitive ratiometric SNARF dye (**Supplementary Figure S16**), we observed that the tau-heparin condensates sustain an internal pH of ∼7.5, slightly more basic compared to the coexisting dilute phase despite the lack of a physical barrier (**Figure 6**a). In contrast to homotypic condensates, heterotypic condensates have a diminished intrinsic pH buffering capacity owing to a reduced thermodynamic need to sustain large proton gradients to maintain stability.^58^ Consequently, the interior pH of tau-heparin coacervates can still scale with solution pH though with a slope less than one (**Figure 6**b). This suggests that the condensed phase makes the internal environment significantly less sensitive to external pH fluctuations within the range studied here.

**Figure 6.**
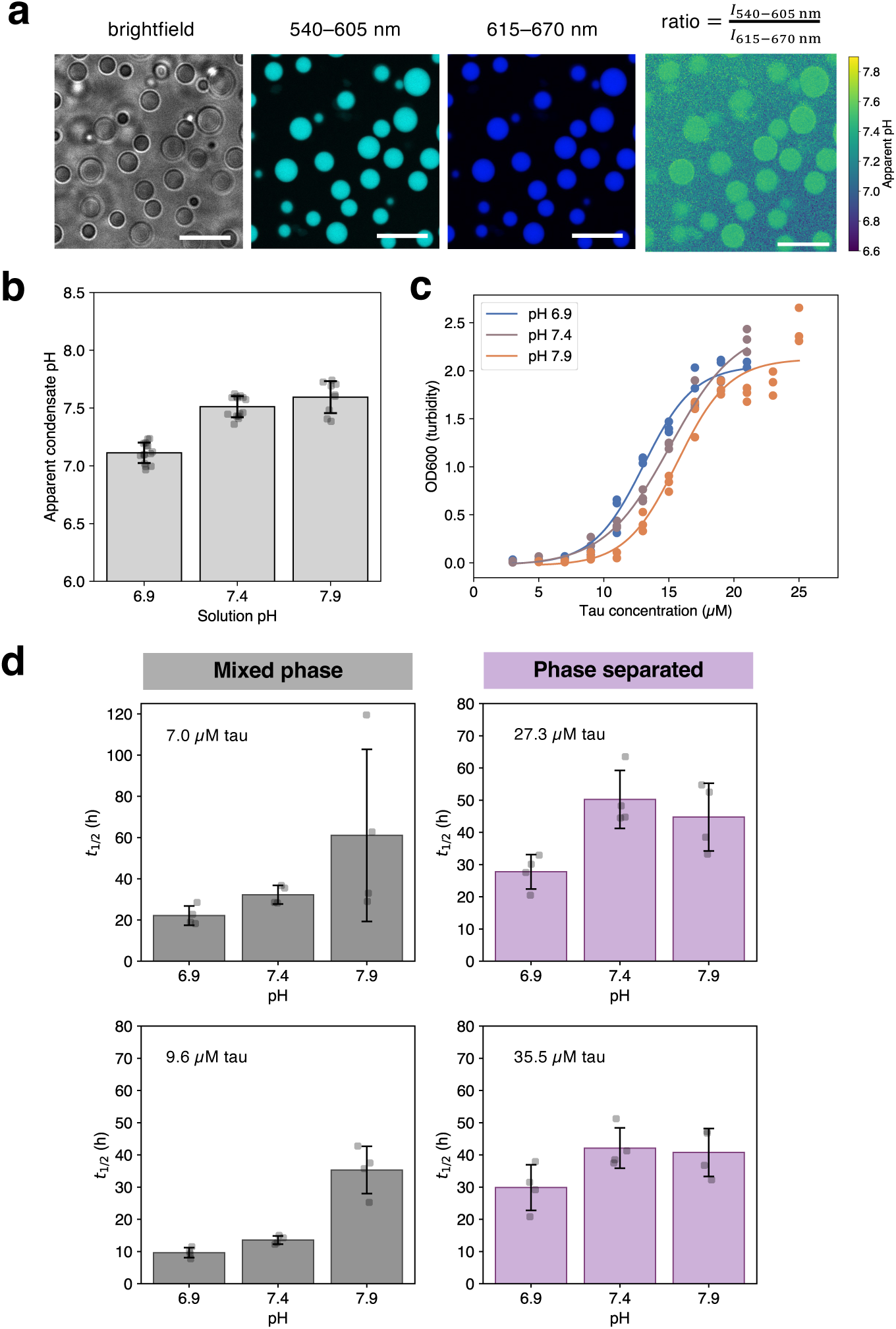
Condensation attenuates the pH sensitivity of aggregation kinetics. (a) Condensates can sustain an internal pH microenvironment. Brightfield, SNARF fluorescence, and ratiometric pH images of tau-heparin condensates at pH 7.4. Scale bars = 10 µm. (b) Apparent condensate pH at different solution pH values. (c) Turbidity measurements illustrating the effect of solution pH on tau-heparin phase behavior at a constant heparin concentration of 80 µg mL^−1^. (n = 3) (d) Effect of solution pH on aggregation half-time *t_1/2_* for mixed-phase vs. phase-separated samples. Aggregation reactions were performed in quadruplicates.

Based on these observations, we hypothesized that the condensate microenvironment could functionally protect tau aggregation from pH changes by buffering the local chemical environment. To test this, we perturbed solution pH across one pH unit while monitoring aggregation kinetics in both the mixed and condensed phases. Our analysis focuses on the phase-specific behavior and response to pH changes. As the binodal boundary shifts with solution pH due to protein protonation changes (**Figure 6**c), we selected specific tau concentrations based on the phase behavior: two concentrations (7.0 µM, 9.6 µM) where tau-heparin mixtures remain homogeneous across all tested pH conditions and two concentrations (27.3 µM, 35.5 µM) where tau becomes compartmentalized within phase-separated droplets.

In the mixed phase, we found that increasing the solution pH over a one-unit range resulted in a pronounced deceleration of aggregation kinetics, characterized by an extended *t*_1/2_ (**Figure 6**d), with normalized ThT curves shown in **Supplementary Figure S17**. This trend is consistent with the electrostatic nature of tau-heparin interactions. Increasing the pH towards the isoelectric point of tau fragment (∼9.9) leads to reduced binding affinity, where the effective local concentration of heparin-bound nucleation centers is reduced, subsequently slowing down amyloid formation.^54,59^

On the other hand, a markedly reduced sensitivity of aggregation kinetics in phase-separated samples to bulk pH perturbations was observed, as evidenced by the fold-change analysis (**Supplementary Figure S18**). We attribute this thermodynamic resilience to the buffering capacity of the condensate microenvironment, which dampens variations in the effective physicochemical conditions experienced by the partitioned tau proteins. As a result, aggregation kinetics within condensates become partly shielded from bulk pH perturbations, effectively suppressing the pH-induced shifts in nucleation kinetics observed in solution. Fibrillation via the liquid-to-amyloid transition thus proceeds under a condensate-defined regime, rendering the aggregation more resilient towards pH fluctuations.

## Discussion

Biological condensates concentrate proteins and other biomolecules through LLPS, and increasing evidence suggests their ability to alter reaction rates. A common view is that condensates accelerate protein aggregation by concentrating it. Yet this concentration-centric perspective overlooks the condensate interior as a distinct reaction microenvironment. We provide evidence that the phase separation does not simply accelerate solution-phase aggregation but alters the tau amyloid pathway. More broadly, this study establishes a tractable model system for probing aggregation within heterotypic condensates, which are prevalent in cellular environments and increasingly implicated in protein misfolding and disease.

Using tau-heparin as a model system, we showed that upon phase separation via complex coacervation, amyloid fibril nucleation and growth are largely restricted to the dense phase. Nucleation becomes largely confined to these dense droplets, which act as highly nucleation-competent environments that substantially lower the classical solution-phase nucleation barrier. Simultaneously, the coexisting dilute phase contributes little to the overall amyloid formation.

This compartmentalization defined by phase equilibrium is the mechanistic basis for the distinct kinetic regime we observed. As total tau concentration increases within the two-phase regime, excess protein partitions predominantly into the dilute phase, while the composition of the coacervate phase remains effectively unchanged within experimental resolution. Consequently, aggregation kinetics are largely decoupled from total protein concentration: increasing total tau does not substantially accelerate nucleation because the reacting population is not the bulk but the fraction sequestered within condensates. This holds true across a variety of tau-polyanion systems, suggesting a general consequence of phase separation.

Parallel to this kinetic divergence, condensation can also alter the structural outcomes of amyloid assembly. Fibrils matured within condensates exhibit mesh-like morphologies that diverge from the long, straight fibrils seen in homogeneous solution. Spectroscopic analyses of amyloid fibrils derived across phase states further highlight that condensation by complex coacervation favors amyloid products with distinct morphological and spectroscopic properties.

Moreover, the condensed phase is not a passive concentrator but an active microreactor that desensitizes aggregation to environmental fluctuations. When challenged with varying solution pH, aggregation kinetics in the mixed phase showed strong sensitivity. Under LLPS, this sensitivity was greatly attenuated. Condensation does not eliminate pH dependence but reduces its dynamic range, providing functional robustness not observed in homogeneous environments. This attenuation reflects the ability of condensates to buffer their internal chemistry, rendering the aggregation process itself more resilient to external perturbations.

Our findings connect closely to recent work showing that homotypic A1-LCD condensates are thermodynamically metastable with respect to fibrils, and that their interiors can act as sinks that suppress, rather than promote, fibril formation.^30^ While this suppression appears to contrast with our observation that tau coacervates are highly nucleation-competent, both behaviors share a common thermodynamic foundation. Above the binodal, the compositions of the coexisting dilute and dense phases are fixed by phase equilibrium. Consequently, increasing the total protein concentration alters the volume fraction of the phases rather than the internal dense-phase reaction environment. Crucially, Das *et al.* show that an increasing dense-phase volume fraction increases the available interfacial area, which actively drives down lag times by accelerating heterogeneous nucleation. The difference in outcome in terms of aggregation suppression versus acceleration may further reflect where each system lies along the metastability axis that they define. Condensates with high sink potential can sequester protein and slow its conversion into fibrils, whereas the electrostatically crosslinked, heterotypic tau-polyanion coacervates studied here favor nucleation. Consistent with both studies, our data shows reduced nucleation barrier for fibrillation in and from condensates. Nucleation at or near the dilute-dense interface^25^, followed by growth that draws on material from the surrounding phases, is compatible with our observations and with interfacial nucleation reported for A1-LCD, FUS, and similar systems.

A weak dependence of aggregation kinetics on total protein concentration within condensates was previously reported for α-synuclein, where it was attributed to concentration buffering of the dense phase by phase equilibrium.^60,61^ Our results extend this principle in three respects. First, they establish it in heterotypic tau-polyanion complex coacervates formed without exogenous crowding agents, where the dense-phase environment is defined by protein-polyanion interactions. Second, they demonstrate that phase state-mediated change in amyloid aggregation kinetics is general across chemically distinct coacervates. Third, they show that, beyond reshaping aggregation kinetics, the condensed state can alter the structure of the resulting amyloid and buffer the aggregation process against pH perturbations. Together with the recent thermodynamic analyses of homotypic condensate metastability mentioned above^30^, these findings point to a common organizing principle: above the binodal, phase equilibrium fixes the composition of the dense phase, while the specific chemistry and material state of the condensate determine whether that environment suppresses or promotes amyloid conversion.

Overall, these findings indicate that the phase state is an important determinant of both the mechanism and the product of tau amyloid formation. Condensation does not merely concentrate tau or accelerate its canonical aggregation pathway; instead, maturation within condensates follows an alternative trajectory, capable of producing fibrils with distinctive structural and kinetic profiles. Moreover, LLPS also reduces the sensitivity of tau aggregation to bulk pH changes, showing that condensates can buffer not only composition but also reaction conditions. These results suggest that condensates modulate pathological protein assembly by setting local reaction conditions, rather than only by increasing reactant concentration. These findings could also inform strategies that target the condensate microenvironment or the liquid-to-amyloid transition.

## Methods

### Materials

Heparin (heparin sodium salt from porcine intestinal mucosa), polyuridylic acid (poly-rU RNA), and poly-L-glutamic acid were purchased from Sigma Aldrich. Fluorescein-conjugated heparin (FITC-heparin) was purchased from Thermo Fisher Scientific. Reagents were used without further purification.

### Tau fragment expression and purification

The K12 tau fragment was expressed and purified using a published protocol.^62^ The construct was engineered by substituting cysteine residues with alanine and appending an N-terminal His₆ tag (K12CFh, cysteine-free, His-tagged). The mutated expression cassettes were synthesized and cloned by GenScript into the pET-28a. The construct was expressed in *Escherichia coli* BL21(DE3). For expression, a single colony was used to inoculate LB medium and grown overnight. Cells were harvested by centrifugation (5,000 rpm, 20 min), and pellets were resuspended in Buffer A (40 mM sodium phosphate, 5 mM imidazole, 400 mM NaCl). Cells were first lysed by sonication, and then was subjected to heat treatment, followed by centrifugation (14,000 × g, 30 min). The supernatant was filtered and loaded onto a HisTrap FF, 5 mL column (Cytiva) pre-equilibrated in Buffer A. Protein was eluted with a step gradient of Buffer B (40 mM sodium phosphate, 300 mM imidazole, 400 mM NaCl) from 10% to 100%. Peak fractions were analyzed by gel electrophoresis (SDS-PAGE), pooled based on purity, and precipitated with acetone overnight. Precipitated protein was collected by centrifugation (4,700 × g, 20 min) and solubilized in 6 M guanidine hydrochloride (GdnHCl). The solubilized pool was further purified by size-exclusion chromatography (SEC) on a HiLoad 26/600 Superdex 75 pg column (Cytiva) equilibrated in Buffer C (15 mM sodium phosphate dibasic, 5 mM sodium phosphate monobasic, 200 mM NaCl). Peak fractions were collected and analyzed by SDS-PAGE. Pure fractions were pooled and lyophilized in aliquots for storage and subsequent use.

### Droplet microfluidics for phase diagram mapping

An in-house combinatorial droplet microfluidic platform termed PhaseScan was used for mapping high-resolution phase diagrams.^38,39^ Polydimethylsiloxane (PDMS) microfluidic devices were fabricated by soft lithography and bonded to glass coverslips. Stock solutions of protein, polyanion, and buffer with separate fluorescent barcodes were dispensed into the device using pressure-driven pumps (Fluigent). Water-in-oil microdroplets were produced via the segmentation of aqueous stream by a surfactant-containing fluorinated oil phase (RAN Biotechnologies). Microdroplets of different compositions were prepared by flowing each component at defined rates, with all aqueous flow rates summing up to 1.4 µL min^−1^. After incubation on-chip, droplets were imaged by multichannel epifluorescence microscopy and analyzed off-line, and biomolecule concentrations were determined using calibration curves generated from labeled solutions of known concentrations.

### Preparation of imaging wells

PDMS slabs (∼4 mm thick) were fabricated using the SYLGARD™ 184 Silicone Elastomer Kit by mixing the polymer base and curing agent at a 10:1 w/w ratio. The mixture was degassed and cured at 65°C. Circular openings, 5 mm in diameter, were made into the cured slabs using a biopsy punch. The slabs were bonded to VWR^®^ cover glass slides (0.13–0.16 mm in thickness) through oxygen plasma activation. Prepared wells were surface-treated with 0.1 w/v% Tween 80 for 30 min, washed with buffer and dried prior to use.

### Confocal microscopy and fluorescence recovery after photobleaching (FRAP)

Confocal laser scanning microscopy was performed on a Leica Stellaris 5 confocal microscope equipped with a 63× oil immersion objective (NA numerical aperture 1.4). Samples containing labeled protein and polyanions in 20 mM HEPES buffer, pH 7.4 were prepared at desired compositions, incubated for 10 min, and imaged in PDMS-based imaging wells (sample volume = 50 µL) at room temperature. Image processing and analysis were performed with FIJI.^63^

FRAP assays were performed on tau fluorescently labeled using Alexa Fluor™ 488 NHS Ester, and photobleaching on condensates was performed using a 488-nm optically pumped semiconductor laser (OPSL). FRAP recovery curves were normalized after correcting for photofading using reference condensates in the same field of view, i.e., 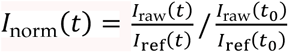.

### Aggregation assay

Lyophilized protein was dissolved in GdnHCl and subsequently monomerized by SEC on a Superdex 75 10/300 GL column (Cytiva) equilibrated in 20 mM HEPES. Peak fractions were collected and used immediately for experiments. Reaction mixtures containing HEPES buffer, monomerized tau, polyanions, and 10 µM ThT were prepared. Samples were dispensed into 384-well optical-bottom plates (Nunc), sealed, and incubated in a plate reader (BMG Labtech) at 25 °C with orbital shaking (60 s, 500 rpm or specified otherwise) alternating with 60 s rest. ThT fluorescence was recorded every 15 min.

Raw intensity traces were screened for aggregation by requiring a >2.5-fold increase over baseline, and positive traces were normalized and smoothed using a Savitzky-Golay filter. Aggregation lag phase duration *t_lag_* was defined as the first time the signal exceeded 0.005 above baseline, and aggregation half-time *t_1/2_* was defined as the first time point at which the smoothed signal reached 0.5.

### Isolation of dilute phases

For a given composition of the phase-separating mixture, >500 µL of sample was prepared. The mixture was centrifuged at 15,000 rpm for 20 min at 4 °C, and the coexisting dilute phase was collected by carefully harvesting the supernatant.

### Quantification of dense phases

Phase-separated samples were prepared as water-in-oil microdroplets stored in PDMS chip. For each composition, dense phase volume fractions were quantified from 3D confocal fluorescence image stack using FIJI. Total droplet volume was modeled as a compressed discoidal geometry with 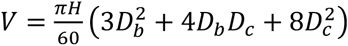, where H is the volume height, D_b_ is the basal plane diameter, and D_c_ is the central plane diameter in µm. The total volume of the dense phase within the same droplet was determined by applying an intensity-based threshold to selectively segment the condensate-rich regions across the z-stack, and dense phase volume fraction was calculated as 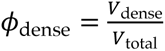. This workflow was benchmarked using an aqueous two-phase system (ATPS) composed of polyethylene glycol and dextran with known volume fraction of phases. Finally, the chemical composition of the dense phase was determined via mass balance constraint:

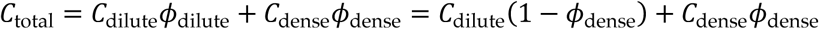

### Transmission electron microscopy

Transmission electron micrographs were acquired on a Talos F200X G2 TEM (Thermo Scientific) operating at 200 keV. Carbon-coated copper grids were glow-discharged prior to use. 2.5 µL of each sample collected from microwell plate was applied to each grid and allowed to adsorb for 40 s, followed by blotting with filter paper to remove excess liquid. Grids were then stained with 2.5 µL of 2 w/v% uranyl acetate for approximately 40 s, blotted, and air-dried at room temperature prior to imaging.

### Dynamic light scattering

The hydrodynamic sizes of heparin-tau complexes and coacervates were determined by dynamic light scattering (DLS) performed on a Zetasizer Nano ZS (Malvern) at room temperature. Each sample was loaded into a quartz cuvette, and the scattering intensity was recorded at a 173° backscattering angle. Three independent repeats were measured.

### Fluorescence spectroscopy

Fluorescence emission spectra were acquired using a Cary Eclipse fluorescence spectrophotometer (Agilent Technologies). Excitation-emission scans were performed by varying the excitation wavelength from 380 to 440 nm in 5-nm increments, and emission spectra over a wavelength range of 450–600 nm were recorded. All measurements were performed at room temperature, with a scan rate of 600 nm min⁻¹ and a spectral resolution of 1 nm per step. Dataset was normalized and smoothed using a Gaussian filter and subsequently interpolated onto a uniform grid (300 × 300 points) to produce 2D excitation-emission maps.

### Attenuated Total Reflectance Fourier Transform Infrared (ATR-FTIR) spectroscopy

Recovered amyloid samples were centrifuged at 15,000 rpm for 30–60 min at 4 °C and pelletized, which were then resuspended in water. ATR-FTIR absorbance spectra were collected on a VERTEX 70 FT-IR Spectrometer (Bruker). 3 µL of the sample was drop-cast and allowed to air-dry. For spectral acquisition, 120 scans were averaged over the range 4,000–900 cm⁻¹ at a resolution of 4 cm⁻¹. Sample absorbance spectra were normalized within the spectral window of interest. Second-derivative plots were calculated and smoothed using a Savitzky-Golay filter, followed by normalization within the respective spectral window with the value at 1800 cm⁻¹ set to one. Spectra and derivative plots were averaged based on multiple repeats.

### Ratiometric pH imaging

Ratiometric pH imaging was performed on the Leica Stellaris 5 confocal microscope with samples spiked with 20 µM SNARF™-4F 5-(and-6)-Carboxylic Acid dye (Thermo Fisher). Samples were excited with a 517-nm laser at 15% power intensity, and the emission intensity in two separate spectral channels, 540–605 nm and 615–670 nm, were collected. Their ratio was used to estimate the solution or condensate pH using a calibration curve, with calibration samples being 20 mM HEPES buffers with different pH values. Ratiometric pH images were generated using Python.

## Author Contributions

Conceptualization: Z.H., P.X.; Investigation: Z.H., P.X., Z.X., Y.W.; Resources: P.X., A.S.; Supervision: T.P.J.K., M.V.; Funding acquisition: T.P.J.K., M.V.; Writing – original draft: Z.H.; Writing – review & editing: Z.H., P.X., M.V., T.P.J.K.

## Acknowledgements

Z.H. thanks Dr. Tianyi Jin (California Institute of Technology) for helpful discussions. We acknowledge funding from UKRI (grants 10059436, 10061100, 10138075, M.V.) and the European Research Council under the European Union’s Seventh Horizon 2020 research and innovation program through the ERC grant DiProPhys (agreement ID 101001615, T.P.J.K.). This work is also supported by EPSRC Underpinning Multi-User Equipment Call (grant number EP/P030467/1).

## References

1. Hampel, H. et al. The Amyloid-β Pathway in Alzheimer’s Disease. Mol. Psychiatry 26, 5481–5503 (2021).

2. Poewe, W. et al. Parkinson disease. Nat. Rev. Dis. Primer 3, 17013 (2017).

3. Knowles, T. P. J., Vendruscolo, M. & Dobson, C. M. The amyloid state and its association with protein misfolding diseases. Nat. Rev. Mol. Cell Biol. 15, 384–396 (2014).

4. Braak, H. & Braak, E. Neuropathological stageing of Alzheimer-related changes. Acta Neuropathol. (Berl.) 82, 239–259 (1991).

5. Jack, C. R. et al. Revised criteria for diagnosis and staging of Alzheimer’s disease: Alzheimer’s Association Workgroup. Alzheimers Dement. 20, 5143–5169 (2024).

6. Lee, V. M.-Y., Goedert, M. & Trojanowski, J. Q. Neurodegenerative Tauopathies. Annu. Rev. Neurosci. 24, 1121–1159 (2001).

7. Wang, Y. & Mandelkow, E. Tau in physiology and pathology. Nat. Rev. Neurosci. 17, 22–35 (2016).

8. Michaels, T. C. T. et al. Amyloid formation as a protein phase transition. Nat. Rev. Phys. 5, 379–397 (2023).

9. Meisl, G. et al. In vivo rate-determining steps of tau seed accumulation in Alzheimer’s disease. Sci. Adv. 7, eabh1448 (2021).

10. Rodriguez Camargo, D. C., et al. Proliferation of Tau 304–380 Fragment Aggregates through Autocatalytic Secondary Nucleation. ACS Chem. Neurosci. 12, 4406–4415 (2021).

11. Santambrogio, A. et al. Serial amplification of tau filaments using Alzheimer’s brain homogenates and C322A or C322S recombinant tau. FEBS Lett. 599, 2768–2778 (2025).

12. Shin, Y. & Brangwynne, C. P. Liquid phase condensation in cell physiology and disease. Science 357, eaaf4382 (2017).

13. Wegmann, S. et al. Tau protein liquid–liquid phase separation can initiate tau aggregation. EMBO J. 37, e98049 (2018).

14. Ray, S. et al. α-Synuclein aggregation nucleates through liquid–liquid phase separation. Nat. Chem. 12, 705–716 (2020).

15. Kanaan, N. M., Hamel, C., Grabinski, T. & Combs, B. Liquid-liquid phase separation induces pathogenic tau conformations in vitro. Nat. Commun. 11, 2809 (2020).

16. Banani, S. F., Lee, H. O., Hyman, A. A. & Rosen, M. K. Biomolecular condensates: organizers of cellular biochemistry. Nat. Rev. Mol. Cell Biol. 18, 285–298 (2017).

17. Ambadipudi, S., Biernat, J., Riedel, D., Mandelkow, E. & Zweckstetter, M. Liquid–liquid phase separation of the microtubule-binding repeats of the Alzheimer-related protein Tau. Nat. Commun. 8, 275 (2017).

18. Alberti, S. & Hyman, A. A. Biomolecular condensates at the nexus of cellular stress, protein aggregation disease and ageing. Nat. Rev. Mol. Cell Biol. 22, 196–213 (2021).

19. Vendruscolo, M. & Fuxreiter, M. Protein condensation diseases: therapeutic opportunities. Nat. Commun. 13, 5550 (2022).

20. Mitrea, D. M., Mittasch, M., Gomes, B. F., Klein, I. A. & Murcko, M. A. Modulating biomolecular condensates: a novel approach to drug discovery. Nat. Rev. Drug Discov. 21, 841–862 (2022).

21. Patel, A. et al. A Liquid-to-Solid Phase Transition of the ALS Protein FUS Accelerated by Disease Mutation. Cell 162, 1066–1077 (2015).

22. Wen, J. et al. Conformational Expansion of Tau in Condensates Promotes Irreversible Aggregation. J. Am. Chem. Soc. 143, 13056–13064 (2021).

23. Šneiderienė, G., et al. Lipid-induced condensate formation from the Alzheimer’s Aβ peptide triggers amyloid aggregation. Proc. Natl. Acad. Sci. 122, e2401307122 (2025).

24. Linsenmeier, M. et al. The interface of condensates of the hnRNPA1 low-complexity domain promotes formation of amyloid fibrils. Nat. Chem. 15, 1340–1349 (2023).

25. Shen, Y., et al. The liquid-to-solid transition of FUS is promoted by the condensate surface. Proc. Natl. Acad. Sci. 120, e2301366120 (2023).

26. Hyman, A. A., Weber, C. A. & Jülicher, F. Liquid-Liquid Phase Separation in Biology. Annu. Rev. Cell Dev. Biol. 30, 39–58 (2014).

27. Boeynaems, S. et al. Protein Phase Separation: A New Phase in Cell Biology. Trends Cell Biol. 28, 420–435 (2018).

28. Visser, B. S., Lipiński, W. P. & Spruijt, E. The role of biomolecular condensates in protein aggregation. Nat. Rev. Chem. 8, 686–700 (2024).

29. Molliex, A. et al. Phase Separation by Low Complexity Domains Promotes Stress Granule Assembly and Drives Pathological Fibrillization. Cell 163, 123–133 (2015).

30. Das, T. et al. Tunable metastability of condensates reconciles their dual roles in amyloid fibril formation. Mol. Cell 85, 2230–2245.e7 (2025).

31. Jawerth, L. et al. Protein condensates as aging Maxwell fluids. Science 370, 1317–1323 (2020).

32. Lipiński, W. P., et al. Biomolecular condensates can both accelerate and suppress aggregation of α-synuclein. Sci. Adv. 8, eabq6495 (2022).

33. Bhandari, K., Sun, Y., Tang, H., Ke, P. C. & Ding, F. A global thermodynamic-kinetic model capturing the hallmarks of liquid-liquid phase separation and amyloid aggregation. Cell Rep. Phys. Sci. 7, 103031 (2026).

34. Metrick, M. A. et al. A single ultrasensitive assay for detection and discrimination of tau aggregates of Alzheimer and Pick diseases. Acta Neuropathol. Commun. 8, 22 (2020).

35. Fichou, Y. et al. Cofactors are essential constituents of stable and seeding-active tau fibrils. Proc. Natl. Acad. Sci. 115, 13234–13239 (2018).

36. Li, D. & Liu, C. Hierarchical chemical determination of amyloid polymorphs in neurodegenerative disease. Nat. Chem. Biol. 17, 237–245 (2021).

37. Falcon, B. et al. Structures of filaments from Pick’s disease reveal a novel tau protein fold. Nature 561, 137–140 (2018).

38. Arter, W. E. et al. Biomolecular condensate phase diagrams with a combinatorial microdroplet platform. Nat. Commun. 13, 7845 (2022).

39. Martens, M. C. M., Han, Z., Knowles, T. P. J., Tuinier, R. & Erkamp, N. A. Protein self-assembly in crowded environments. Preprint at 10.26434/chemrxiv-2025-zhb2l (2025).

40. Banerjee, P. R., Milin, A. N., Moosa, M. M., Onuchic, P. L. & Deniz, A. A. Reentrant Phase Transition Drives Dynamic Substructure Formation in Ribonucleoprotein Droplets. Angew. Chem. Int. Ed. 56, 11354–11359 (2017).

41. R. Tejedor, A., et al. Chemically Informed Coarse-Graining of Electrostatic Forces in Charge-Rich Biomolecular Condensates. ACS Cent. Sci. 11, 302–321 (2025).

42. Makasewicz, K., Morelli, C., Guida, T., Faltova, L. & Arosio, P. Competition between protein-RNA clustering and phase separation drives re-entrant phase behavior of hnRNPA1. Nat. Commun. 10.1038/s41467-026-71939-2 (2026) doi:10.1038/s41467-026-71939-2.

43. Arosio, P., Knowles, T. P. J. & Linse, S. On the lag phase in amyloid fibril formation. Phys. Chem. Chem. Phys. 17, 7606–7618 (2015).

44. Meisl, G. et al. Molecular mechanisms of protein aggregation from global fitting of kinetic models. Nat. Protoc. 11, 252–272 (2016).

45. Cohen, S. I. A. et al. Proliferation of amyloid-β42 aggregates occurs through a secondary nucleation mechanism. Proc. Natl. Acad. Sci. 110, 9758–9763 (2013).

46. Brelstaff, J. et al. The fluorescent pentameric oligothiophene pFTAA identifies filamentous tau in live neurons cultured from adult P301S tau mice. Front. Neurosci. 9, (2015).

47. Zhang, P., Alsaifi, N. M., Wu, J. & Wang, Z.-G. Polyelectrolyte complex coacervation: Effects of concentration asymmetry. J. Chem. Phys. 149, 163303 (2018).

48. Wang, J., Cohen Stuart, M. A. & Van Der Gucht, J. Phase Diagram of Coacervate Complexes Containing Reversible Coordination Structures. Macromolecules 45, 8903–8909 (2012).

49. Fuxreiter, M. & Vendruscolo, M. Generic nature of the condensed states of proteins. Nat. Cell Biol. 23, 587–594 (2021).

50. Sarroukh, R., Goormaghtigh, E., Ruysschaert, J.-M. & Raussens, V. ATR-FTIR: A “rejuvenated” tool to investigate amyloid proteins. Biochim. Biophys. Acta BBA - Biomembr. 1828, 2328–2338 (2013).

51. Biancalana, M. & Koide, S. Molecular mechanism of Thioflavin-T binding to amyloid fibrils. Biochim. Biophys. Acta BBA - Proteins Proteomics 1804, 1405–1412 (2010).

52. Montgomery, K. M. et al. Chemical Features of Polyanions Modulate Tau Aggregation and Conformational States. J. Am. Chem. Soc. 145, 3926–3936 (2023).

53. Zhang, X. et al. RNA stores tau reversibly in complex coacervates. PLOS Biol. 15, e2002183 (2017).

54. Friedhoff, P., Schneider, A., Mandelkow, E.-M. & Mandelkow, E. Rapid Assembly of Alzheimer-like Paired Helical Filaments from Microtubule-Associated Protein Tau Monitored by Fluorescence in Solution. Biochemistry 37, 10223–10230 (1998).

55. Jeganathan, S., Von Bergen, M., Mandelkow, E.-M. & Mandelkow, E. The Natively Unfolded Character of Tau and Its Aggregation to Alzheimer-like Paired Helical Filaments. Biochemistry 47, 10526–10539 (2008).

56. King, M. R. et al. Macromolecular condensation organizes nucleolar sub-phases to set up a pH gradient. Cell 187, 1889–1906.e24 (2024).

57. Dai, Y., Wang, Z.-G. & Zare, R. N. Unlocking the electrochemical functions of biomolecular condensates. Nat. Chem. Biol. 20, 1420–1433 (2024).

58. Ausserwöger, H. et al. Biomolecular condensates sustain pH gradients at equilibrium through charge neutralization. Nat. Chem. 18, 246–257 (2026).

59. Charafeddine, R. A. et al. Tau repeat regions contain conserved histidine residues that modulate microtubule-binding in response to changes in pH. J. Biol. Chem. 294, 8779–8790 (2019).

60. Dada, S. T., et al. Spontaneous nucleation and fast aggregate-dependent proliferation of α-synuclein aggregates within liquid condensates at neutral pH. Proc. Natl. Acad. Sci. 120, e2208792120 (2023).

61. Klosin, A. et al. Phase separation provides a mechanism to reduce noise in cells. Science 367, 464–468 (2020).

62. Santambrogio, A. et al. Design of Tau Aggregation Inhibitors Using Iterative Machine Learning and a Polymorph-Specific Brain-Seeded Fibril Amplification Assay. J. Am. Chem. Soc. 147, 35942–35952 (2025).

63. Schindelin, J., et al. Fiji: an open-source platform for biological-image analysis. Nat. Methods 9, 676–682 (2012).

