## Supplementary material for "Complex coacervation reshapes the aggregation landscape of tau": SI file

Zexiang Han *et al.*

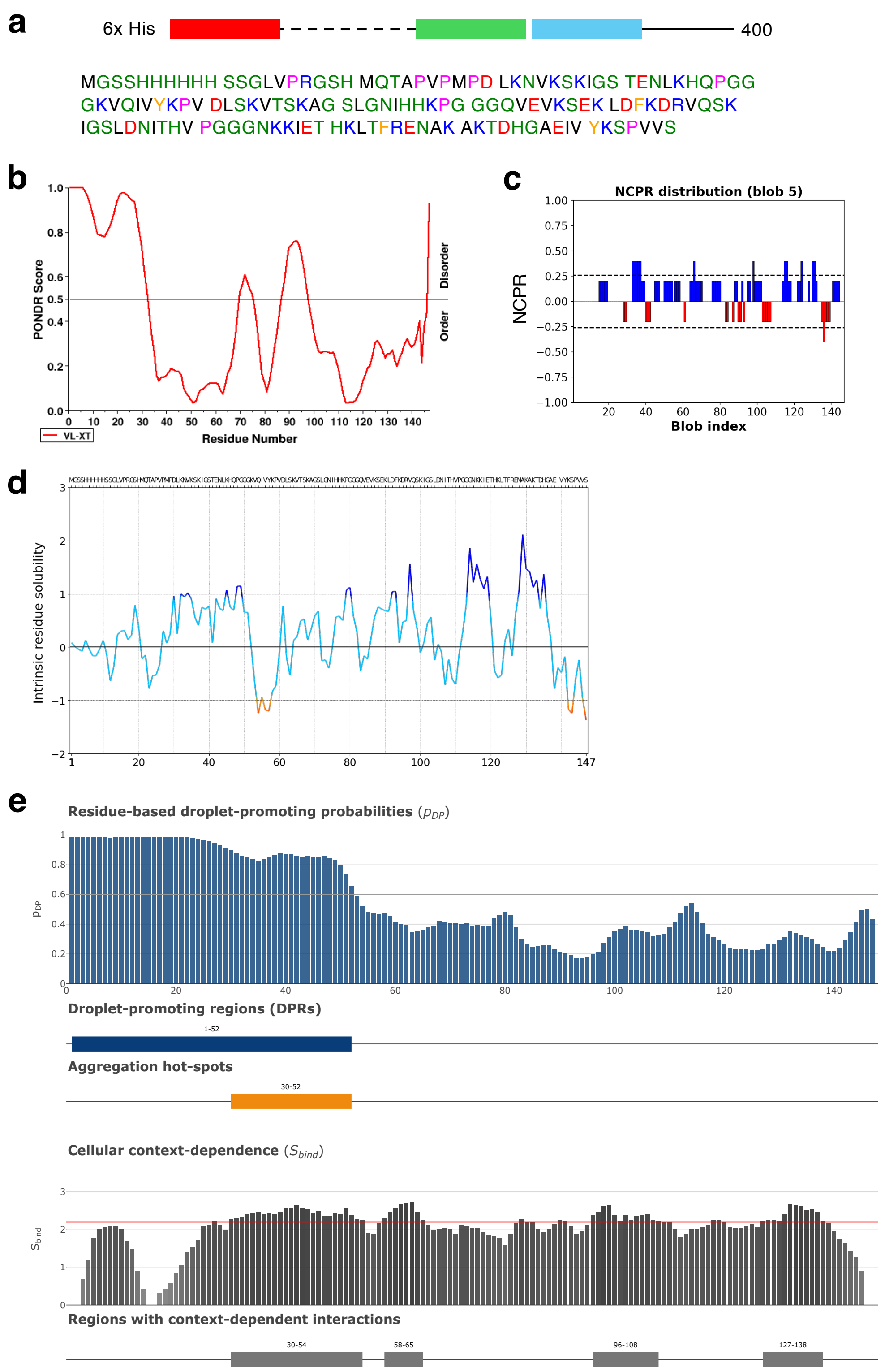


**Supplementary Figure S1. Sequence analysis of the tau fragment used in this work.** (a) Scheme of the K12 construct and its sequence. (b) Order-disorder PONDR score[1]. (c) Charge distribution analyzed using CIDER[2]. (d) Intrinsic solubility profile calculated using CamSol[3]. Here, a value below -1 is aggregation-promoting, above 1 solubility-promoting. (e) Sequence-based prediction of the phase separation propensity using FuzDrop[4]. Droplet-promoting regions (DPRs) identify contiguous sequence segments predicted to drive condensate formation. Aggregation hotspots denote residues with an increased propensity to undergo a transition from dynamic condensate interactions to ordered intermolecular aggregation.

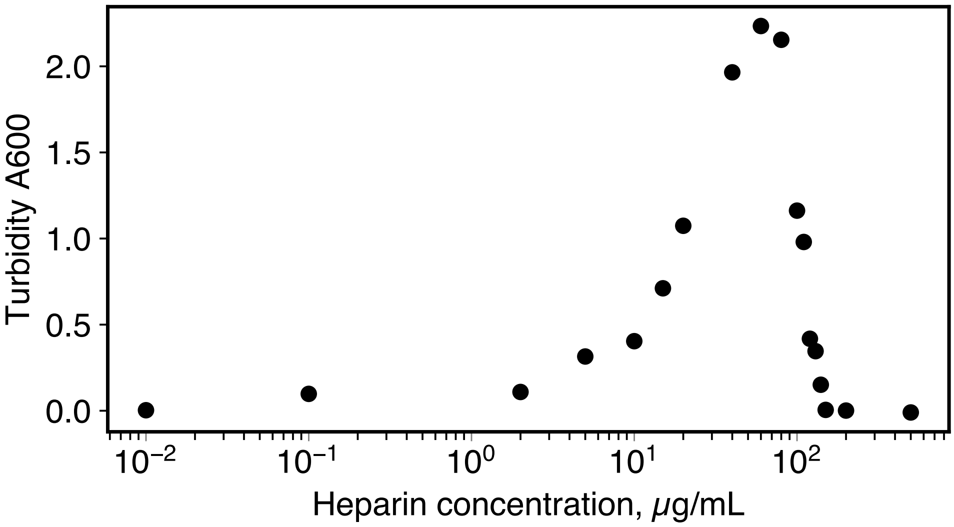


**Supplementary Figure S2. Re-entrant phase behavior of tau-heparin mixtures monitored by turbidity.** Turbidity at 600 nm was measured for tau-heparin mixtures containing 20 $\mu$M tau and increasing concentrations of heparin. The increase in turbidity at intermediate heparin concentrations indicates condensate formation, whereas the subsequent decrease at higher heparin concentrations indicates condensate dissolution, consistent with re-entrant complex coacervation driven by charge and valency balance.

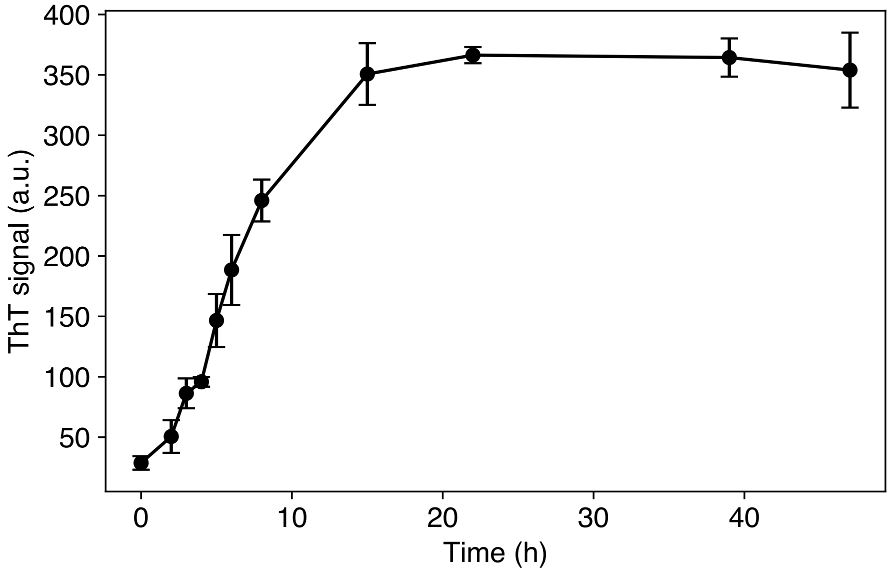


**Supplementary Figure S3. Accumulation of ThT-positive amyloid signal within tau condensates.** The phase-separated mixture contained 35.5 $\mu$M tau and 80 $\mu$g mL$^{-1}$ heparin. The intra-condensate ThT signal increases during early aggregation and then plateaus, indicating that the condensate volume has become saturated with ThT-binding β-sheet structures. The continued increase in bulk ThT signal beyond this plateau therefore reflects an increasing fraction of ThT-positive fibrillar aggregates within the system.

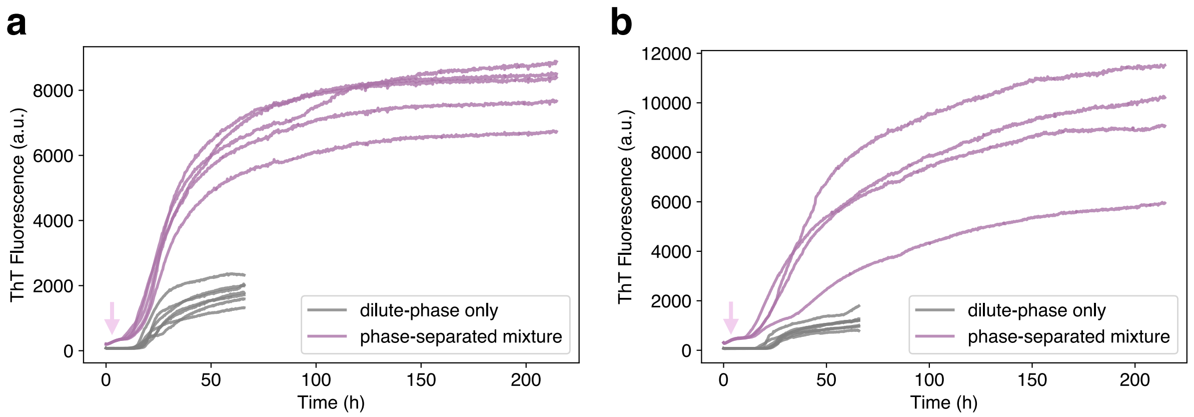


**Supplementary Figure S4. ThT aggregation curves of two phase-separated mixtures versus their isolated dilute phases.** Here, although the ThT traces followed a similar kinetic profile under high shear, the signal intensity was significantly lower than that of the phase-separated ensemble. Pink arrows highlight spontaneous nucleation in the dense phase.

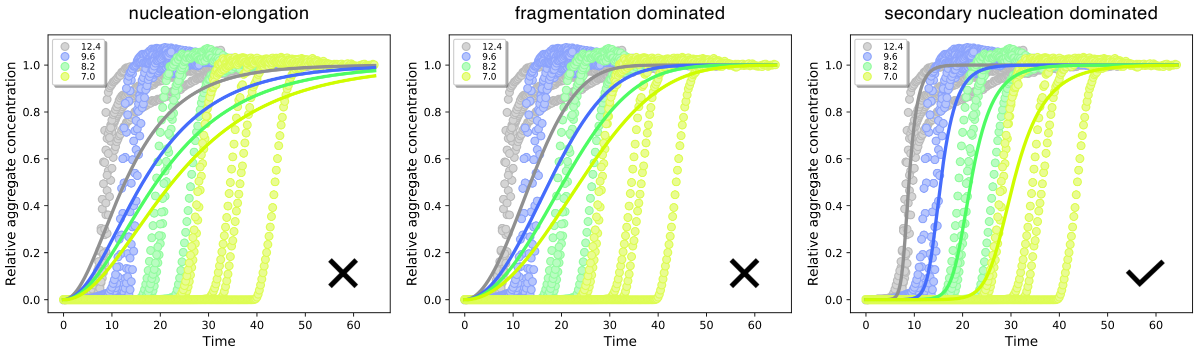


**Supplementary Figure S5. Global kinetic fitting supports secondary nucleation as the dominant mechanism in the mixed phase.** Normalized aggregation kinetics for a dilution series of tau in homogeneous (mixed-phase) solution were fitted globally in AmyloFit using three candidate models: nucleation-elongation (left), fragmentation-dominated (centre), and secondary-nucleation-dominated (right). Points are experimental data and solid lines are the global fits, with shared rate constants across all monomer concentrations. Only the secondary-nucleation-dominated model reproduces the concentration dependence across the full series; the nucleation-elongation and fragmentation models fail to capture it. This assignment is consistent with the steep concentration scaling of the aggregation half-time (apparent exponent -3, see Supplementary Figure S6). Monomer concentrations are given in $\mu$M.

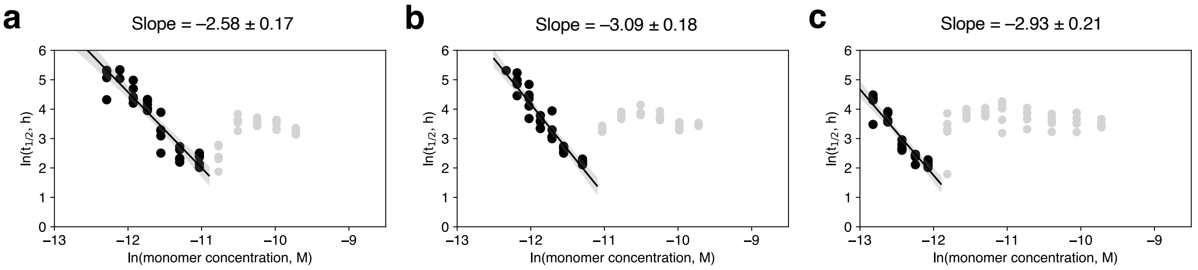


**Supplementary Figure S6. The scaling exponents of the aggregation half-time reveal a strong monomer-concentration dependence in the mixed phase that collapses upon phase separation.** (a-c) Double-logarithmic plots of aggregation half-time against monomer concentration for tau in homogeneous solution at (a) 40, (b) 80 and (c) 120 $\mu$g mL$^{-1}$ heparin. Black points correspond to the mixed (homogeneous) regime; the solid line is a linear fit whose slope gives the apparent scaling exponent (a, -2.58 ± 0.17; b, -3.09 ± 0.18; c, -2.93 ± 0.21). The steep negative exponents (about -3) indicate a strong dependence of aggregation kinetics on monomer concentration, characteristic of a nucleation-and-growth process dominated by secondary nucleation (see Supplementary Figure S5). Grey points lie above the binodal, in the phase-separated regime, which were excluded from the fit and show the loss of concentration dependence that defines the compartmentalized regime analyzed in Figure 4.

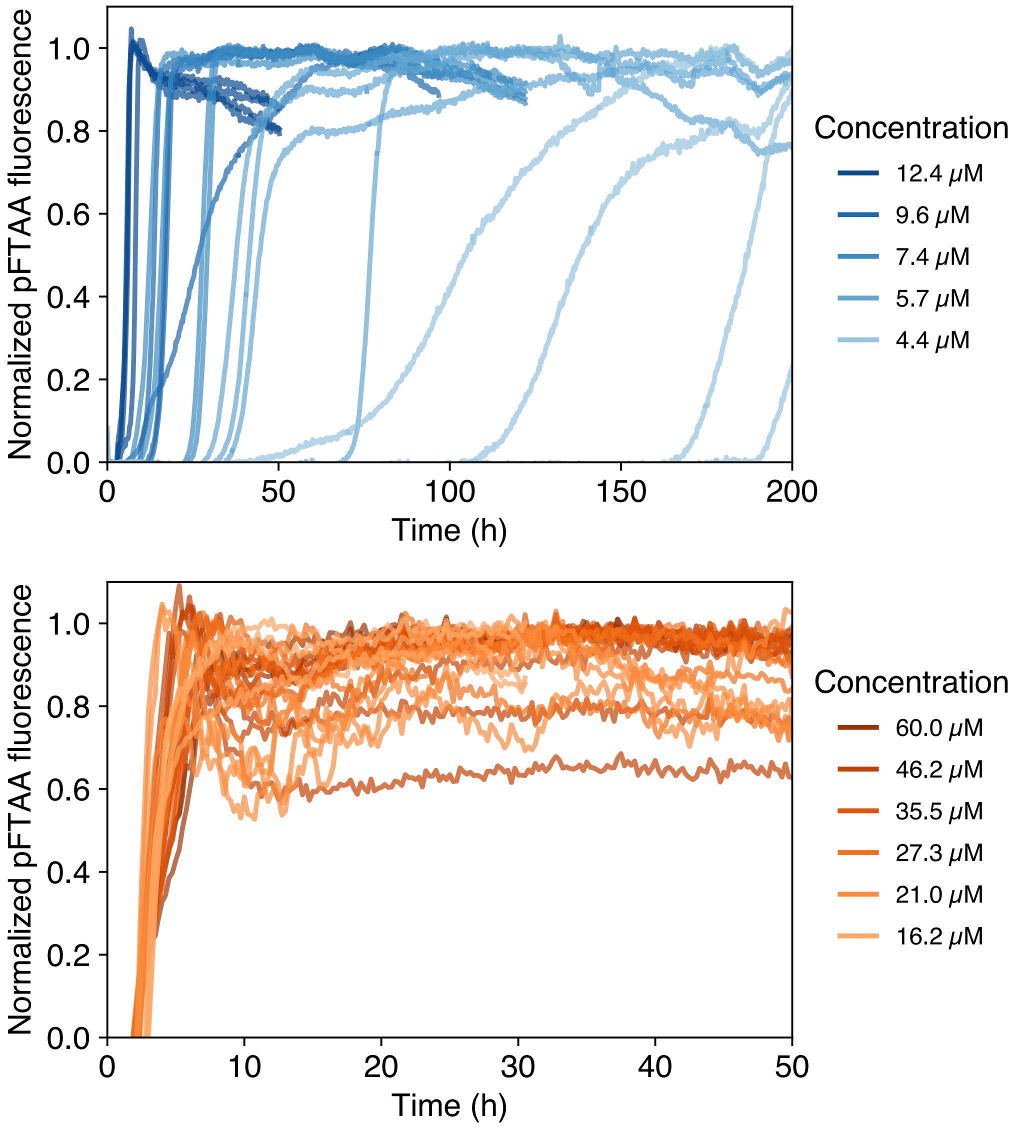


**Supplementary Figure S7. pFTAA reports rapid early-stage assembly that precedes the ThT-detected fibril signal.** Normalized pFTAA fluorescence kinetics for a dilution series of tau in the presence of 80 $\mu$g mL$^{-1}$ heparin, shown for (top) the mixed (homogeneous) regime and (bottom) the phase-separated regime, with monomer concentration indicated by color. pFTAA, an oligothiophene that detects early-stage prefibrillar intermediates, reaches a plateau within the first few hours in phase-separated mixtures, well before the rise of the ThT signal under matched conditions (see Figure 4c), which is consistent with condensates accelerating the initial nucleation step and accumulating prefibrillar species ahead of their slower conversion into canonical, ThT-positive amyloid.

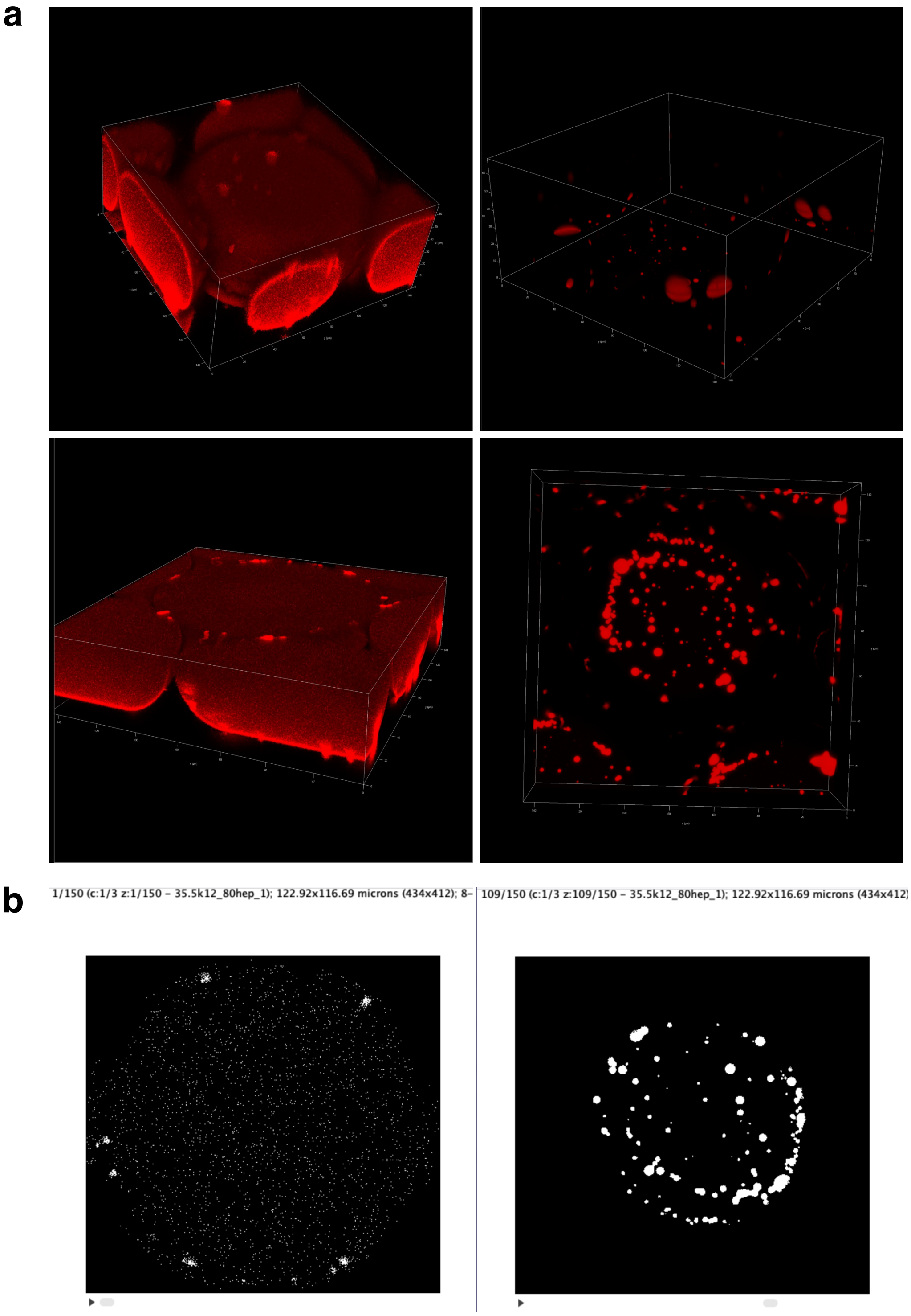


**Supplementary Figure S8. Quantification of volume fractions in microdroplets using confocal z-stacks.** (a) 3D z-stacks showing compressed microdroplets into a discoidal geometry. Condensates were seen mostly at the bottom of the droplet. (b) Selected views of a single microdroplet. Right image shows segmented condensate region used for volume fraction quantification.

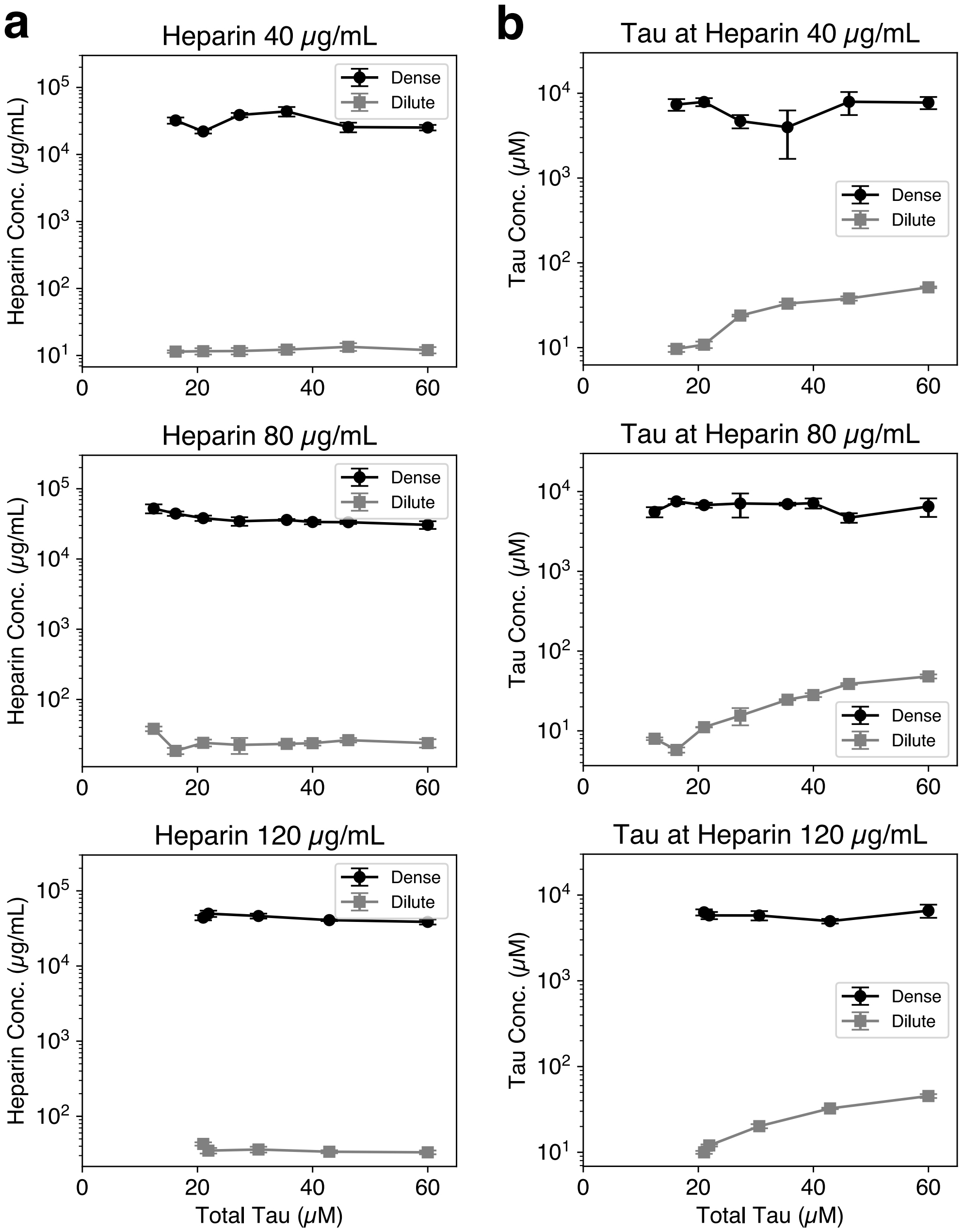


**Supplementary Figure S9. The dense-phase composition is buffered as total tau increases, with excess protein partitioning into the dilute phase.** Estimated dense- and dilute-phase concentrations as a function of total tau concentration, determined from fluorescence measurements combined with the mass-balance and volume-fraction analysis described in the Methods (n = 3; error bars, s.d.). (a) Heparin and (b) tau concentrations in the coexisting dense (filled) and dilute (open) phases at constant total heparin of (top) 40, (middle) 80 and (bottom) 120 $\mu$g mL$^{-1}$ (note the logarithmic concentration axes). Across all three conditions, the dense-phase concentration of both components remains approximately constant over a broad range of total tau, whereas the dilute-phase tau concentration rises with increasing total tau. This invariance of dense-phase composition, which is set by the tau–heparin coacervation equilibrium, is the basis for the weak dependence of aggregation kinetics on total tau concentration reported in Figure 4 and underlies the concentration-buffering mechanism.

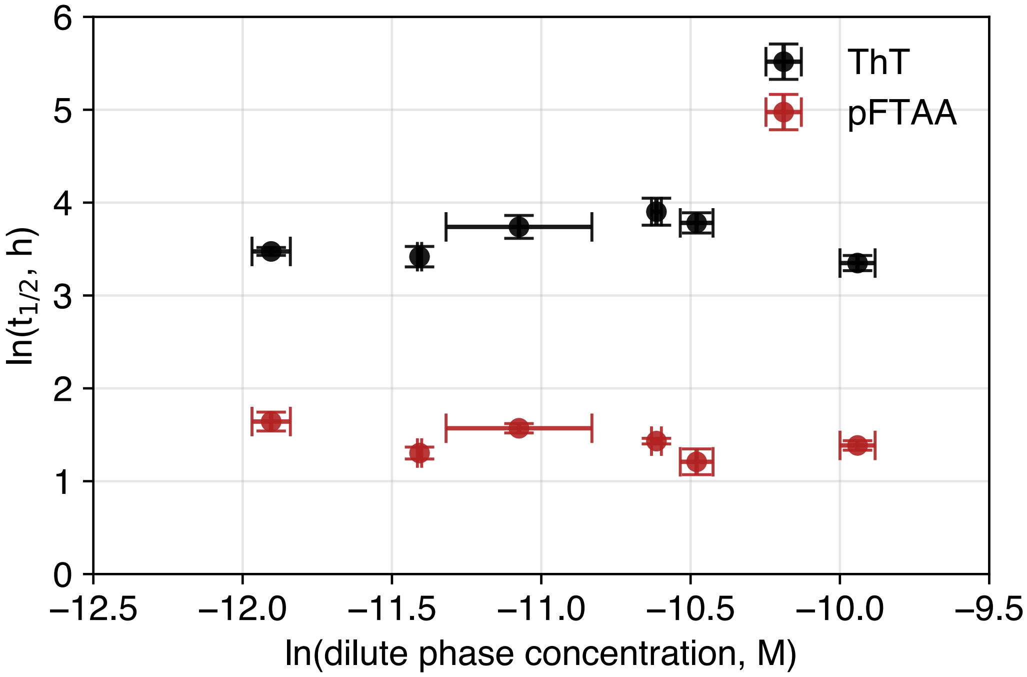


**Supplementary Figure S10. Aggregation kinetics do not track the dilute-phase tau concentration in the phase-separated regime.** Aggregation half-time plotted against the coexisting dilute-phase tau concentration for phase-separated tau-heparin mixtures. Despite a substantial increase in dilute-phase tau concentration with increasing total tau (Supplementary Figure S9), the half-time shows no systematic dependence on it, indicating that the dilute-phase reservoir does not measurably contribute to the aggregation pathway. Together with the near-invariant dense-phase composition (Supplementary Figure S9), this identifies the buffered dense phase, not the dilute phase, as the kinetically relevant compartment, consistent with the concentration-buffering mechanism in Figure 4.

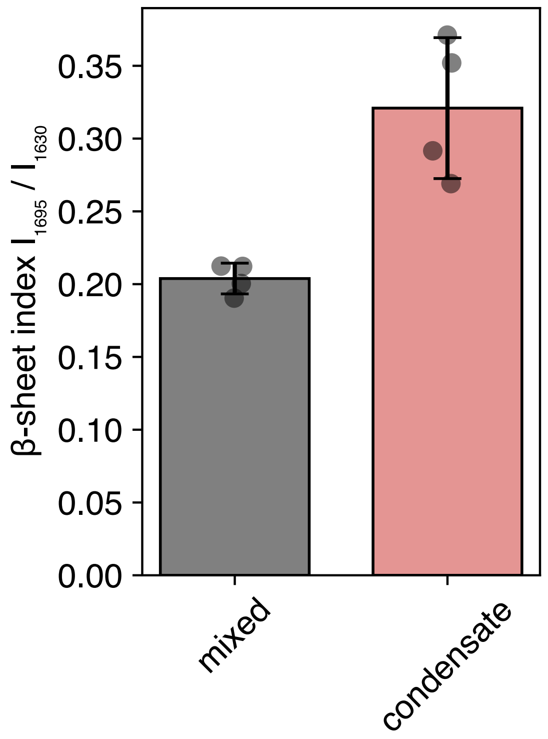


**Supplementary Figure S11. Condensate-derived fibrils show greater antiparallel β-sheet content.** β-sheet index, defined as the ratio of the FTIR second-derivative amide-I intensities at 1695 and 1630 $cm^{-1}$ (a proxy for antiparallel-to-parallel β-sheet organization), for fibrils grown in mixed solution versus within condensates. Bars are mean ± s.d.; points are individual replicates.

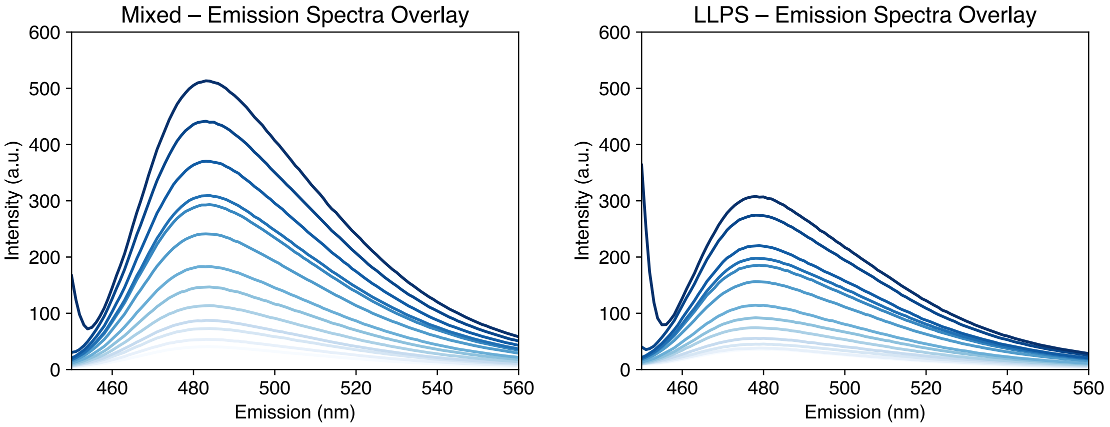


**Supplementary Figure S12. ThT emission depends on fibril phase-of-origin.** ThT emission spectra recorded across excitation wavelengths (380-440 nm) for fibrils grown in mixed solution (left) versus within condensates (right); color denotes excitation wavelength. These spectra are the basis for the 2D excitation-emission maps in Figure 5c, where condensate-derived fibrils show a blue-shifted emission maximum.

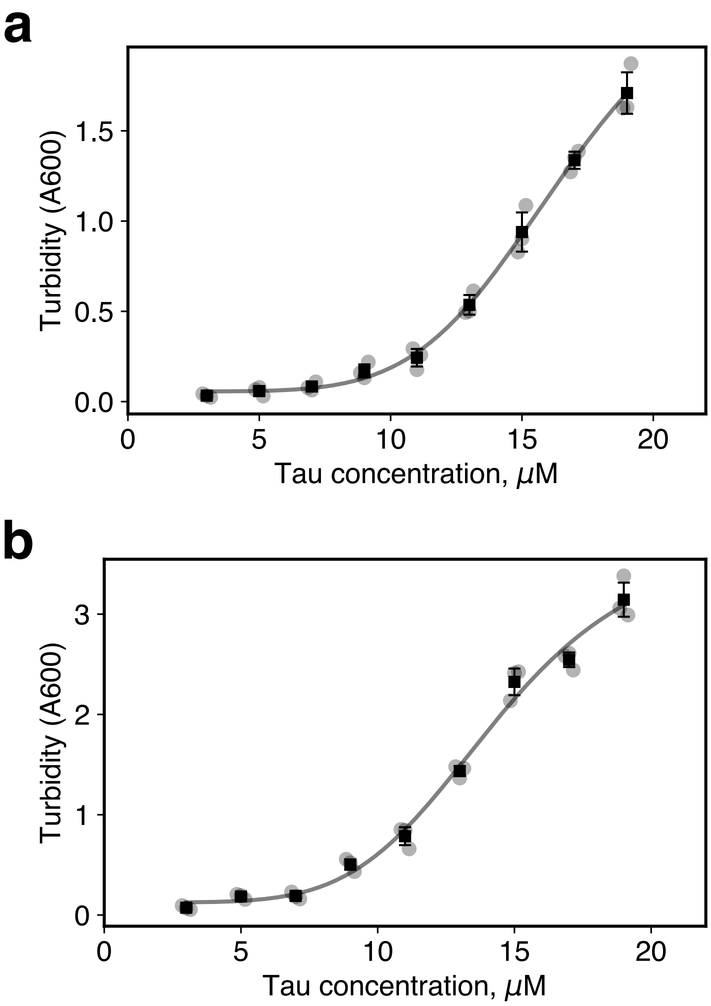


**Supplementary Figure S13. Turbidity measurements illustrating tau-polyanion phase behavior.** Two polyanion concentrations were used: (a) 270 $\mu$g mL$^{-1}$ poly-rU RNA and (b) 100 $\mu$g mL$^{-1}$ poly-L-Glu.

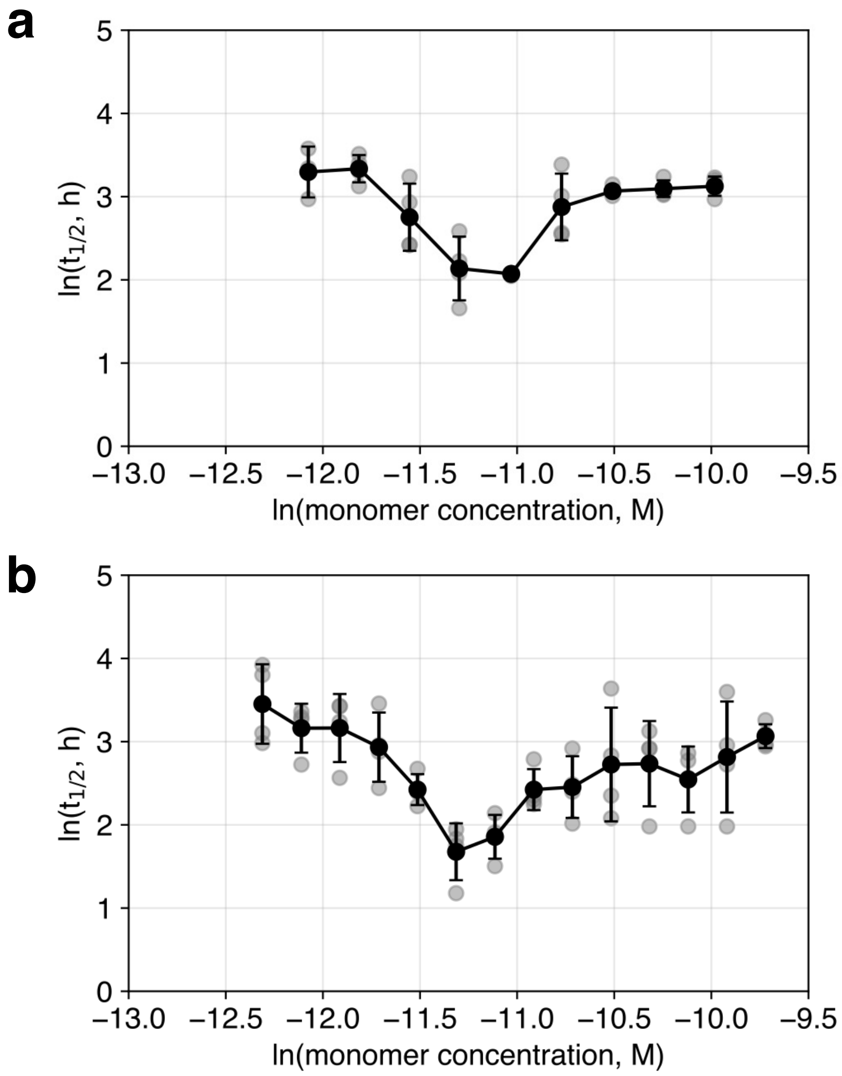


**Supplementary Figure S14. Relationship between tau monomer concentration and aggregation half-time $t_{1/2}$ as monitored by ThT:** (a) for 270 $\mu$g mL$^{-1}$ poly-rU RNA and (b) for 100 $\mu$g mL$^{-1}$ poly-L-Glu.

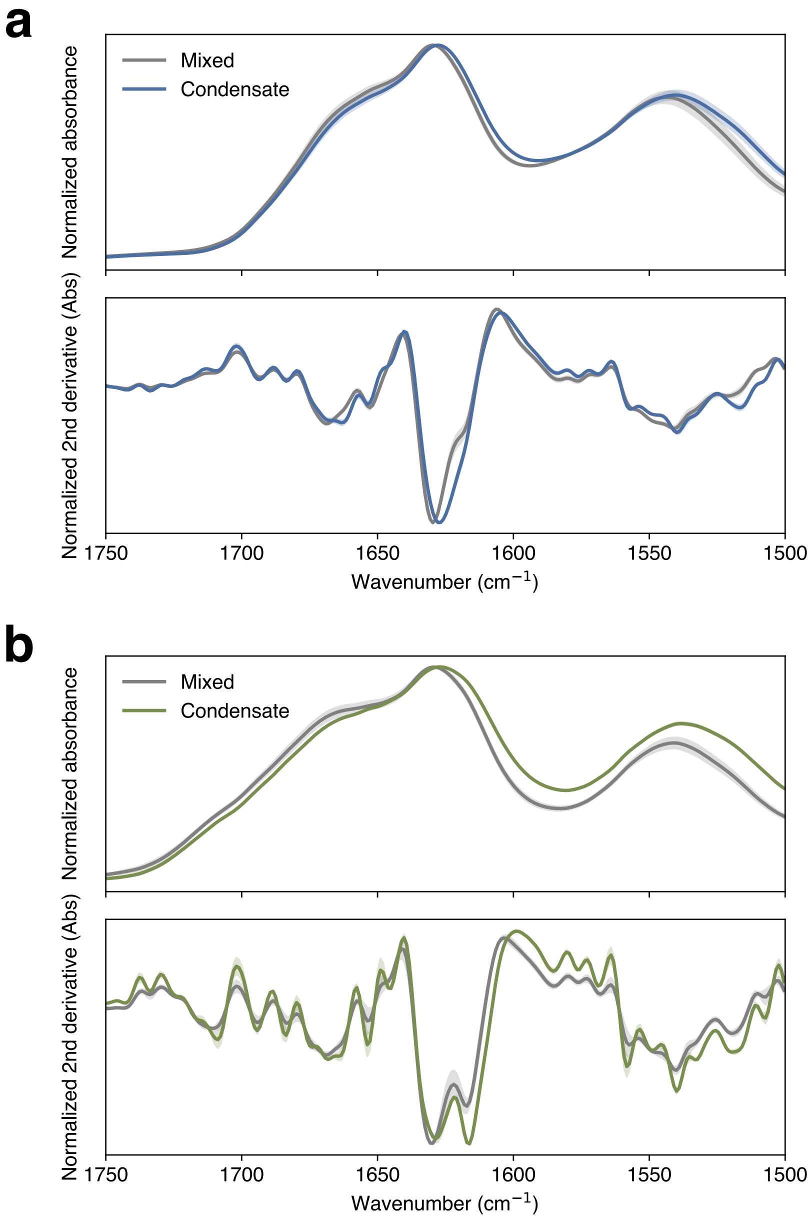


**Supplementary Figure S15. Fibrils formed within condensates of other polyanions show detectable but more modest secondary-structure differences than the heparin system.** Averaged ATR-FTIR absorbance spectra (top of each panel) and second-derivative plots (bottom) over the amide-I region for tau fibrils grown in the mixed phase versus within condensates, for (a) tau-poly-L-glutamate and (b) tau-poly-rU RNA. Error bands show s.d. In both systems the mixed and condensate spectra differ in the amide-I band, indicating phase-state-dependent fibril structure, but the differences are smaller than those seen for tau–heparin (Figure 5b), which is consistent with structural polymorph selection remaining sensitive to polyanion chemistry, even though the concentration dependence of the kinetics is phase-state-dependent across all three systems.

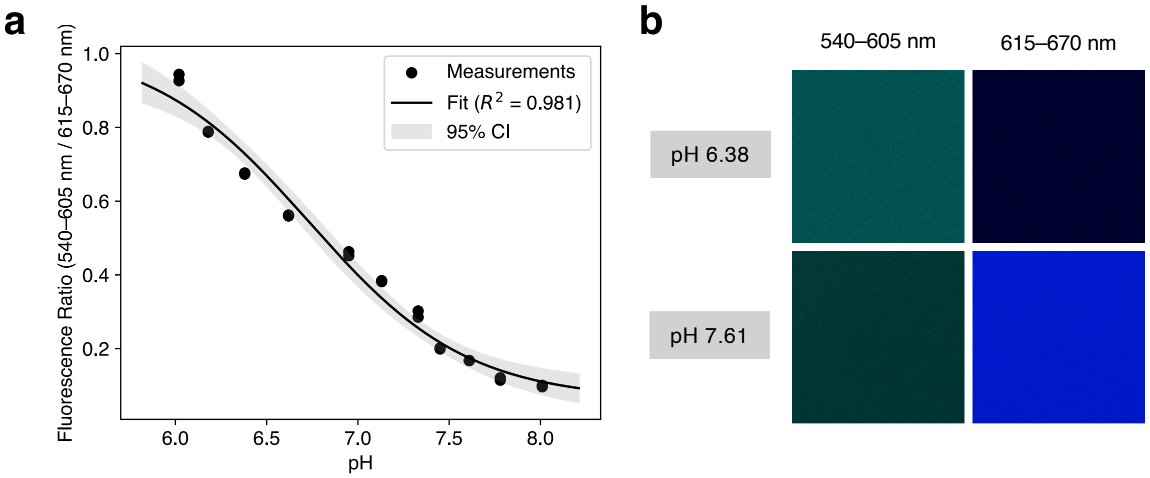


**Supplementary Figure S16. SNARF calibration for ratiometric pH measurement.** (a) Calibration curve relating the SNARF emission ratio (540-605 nm/615-670 nm) to pH, measured in 20 mM HEPES buffers of defined pH; points are measurements and the solid line is the fit ($R^2$ = 0.981) with 95% confidence band. (b, c) Representative confocal images of calibration solutions in the two emission channels at low and high pH. This calibration was used to convert intra- and extra-condensate SNARF ratios into apparent pH values in Figure 6a, b.

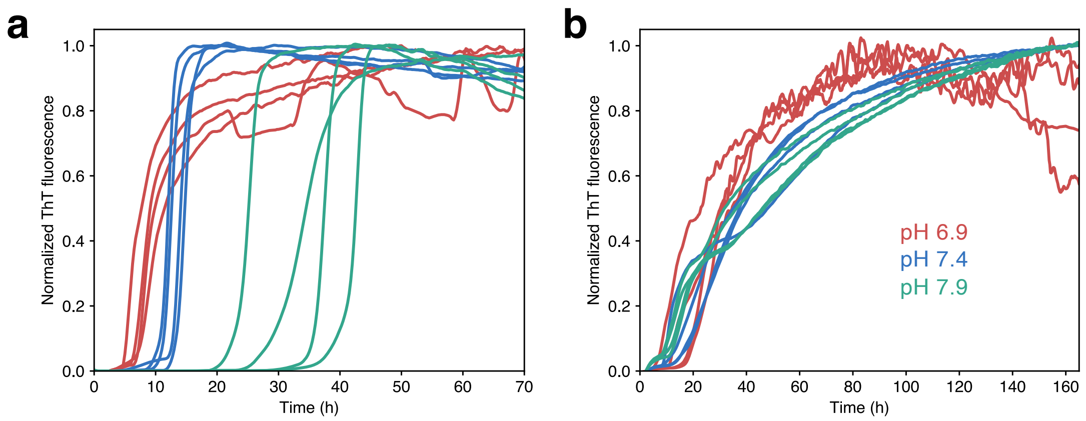


**Supplementary Figure S17. Phase separation attenuates the pH sensitivity of tau aggregation kinetics.** Normalized ThT aggregation traces at solution pH 6.9, 7.4 and 7.9 (color-coded) for tau-heparin mixtures in (a) the homogeneous regime (9.6 $\mu$M tau) and (b) the phase-separated regime (35.5 $\mu$M tau), both at 80 $\mu$g mL$^{-1}$ heparin. In the mixed phase, raising pH markedly slows aggregation (right-shifted curves); in the phase-separated regime this pH dependence is strongly reduced. These traces underlie the half-time comparison in Figure 6d.

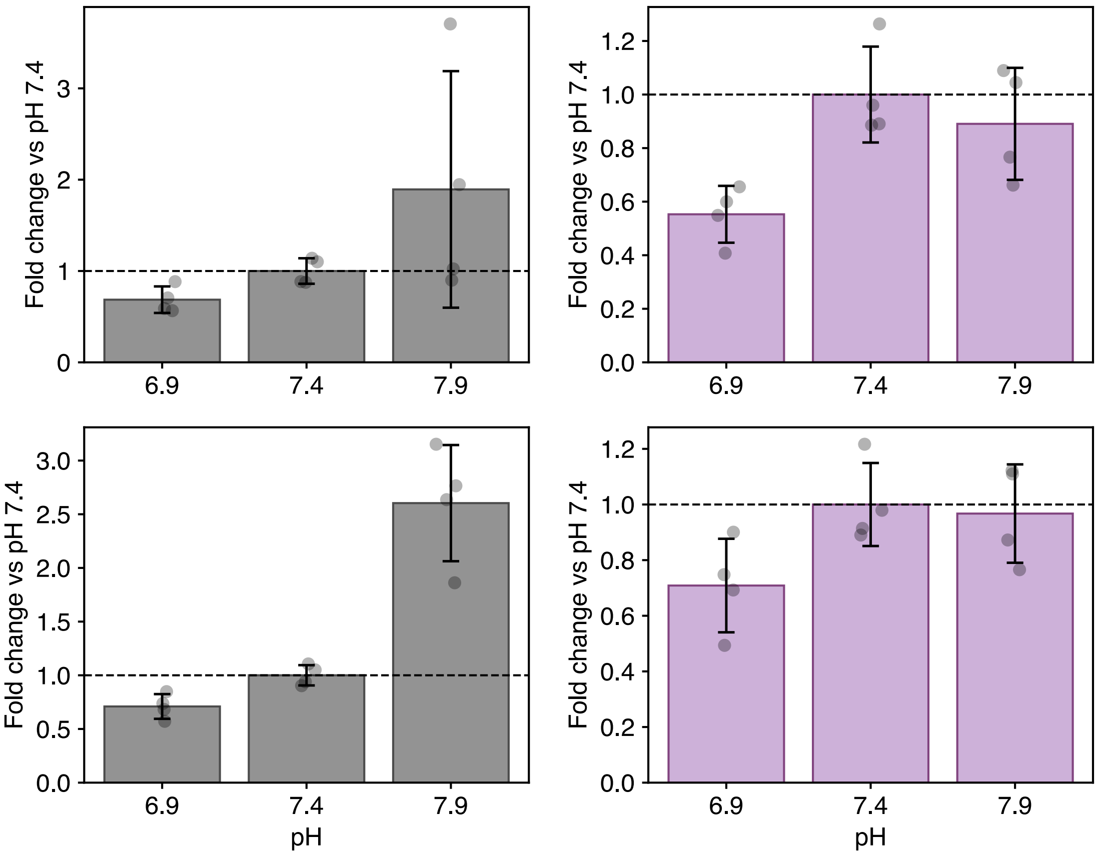


**Supplementary Figure S18. Phase separation attenuates the pH dependence of tau aggregation kinetics.** Aggregation half-times from Figure 6d are shown as fold changes relative to the corresponding value at pH 7.4 for each tau concentration and phase state. Homogeneous samples exhibit larger pH-dependent changes in aggregation kinetics, whereas phase-separated samples show a reduced dynamic range, consistent with partial buffering of the aggregation reaction by the condensate microenvironment.

**Supplementary References**


1. Xue, B., Dunbrack, R. L., Williams, R. W., Dunker, A. K. & Uversky, V. N. PONDR-FIT: A meta-predictor of intrinsically disordered amino acids. *Biochim. Biophys. Acta BBA - Proteins Proteomics* **1804**, 996–1010 (2010).
2. Holehouse, A. S., Das, R. K., Ahad, J. N., Richardson, M. O. G. & Pappu, R. V. CIDER: Resources to Analyze Sequence-Ensemble Relationships of Intrinsically Disordered Proteins. *Biophys. J.* **112**, 16–21 (2017).
3. Sormanni, P., Aprile, F. A. & Vendruscolo, M. The CamSol Method of Rational Design of Protein Mutants with Enhanced Solubility. *J. Mol. Biol.* **427**, 478–490 (2015).
4. Vendruscolo, M. & Fuxreiter, M. FuzDrop: sequence-based prediction of the propensity of proteins for liquid–liquid phase separation and aggregation. *Nat. Protoc.* **21**, 2016–2042 (2026).
